# Accurate delineation of individual tree crowns in tropical forests from aerial RGB imagery using Mask R-CNN

**DOI:** 10.1101/2022.07.10.499480

**Authors:** James G. C. Ball, Sebastian H. M. Hickman, Tobias D. Jackson, Xian Jing Koay, James Hirst, William Jay, Matthew Archer, Mélaine Aubry-Kientz, Grégoire Vincent, David A. Coomes

## Abstract

Tropical forests are a major component of the global carbon cycle and home to two-thirds of terrestrial species. Upper-canopy trees store the majority of forest carbon and can be vulnerable to drought events and storms. Monitoring their growth and mortality is essential to understanding forest resilience to climate change, but in the context of forest carbon storage, large trees are underrepresented in traditional field surveys, so estimates are poorly constrained. Aerial photographs provide spectral and textural information to discriminate between tree crowns in diverse, complex tropical canopies, potentially opening the door to landscape monitoring of large trees. Here we describe a new deep convolutional neural network method, *Detectree2*, which builds on the Mask R-CNN computer vision framework to recognise the irregular edges of individual tree crowns from airborne RGB imagery. We trained and evaluated this model with 3,797 manually delineated tree crowns at three sites in Malaysian Borneo and one site in French Guiana. As an example application, we combined the delineations with repeat lidar surveys (taken between 3 and 6 years apart) of the four sites to estimate the growth and mortality of upper-canopy trees. *Detectree2* delineated 65,000 upper-canopy trees across 14 km^2^ of aerial images. The skill of the automatic method in delineating unseen test trees was good (F_1_ score = 0.64) and for the tallest category of trees was excellent (F_1_ score = 0.74). As predicted from previous field studies, we found that growth rate declined with tree height and tall trees had higher mortality rates than intermediate-size trees. Our approach demonstrates that deep learning methods can automatically segment trees in widely accessible RGB imagery. This tool (provided as an open-source Python package) has many potential applications in forest ecology and conservation, from estimating carbon stocks to monitoring forest phenology and restoration.

**Python package available to install at** https://github.com/PatBall1/Detectree2

## Introduction

Intact tropical forests are an important component of the global carbon cycle: they are major carbon stores and significant carbon sinks (1). However, the strength of the carbon sink is diminishing as a result of global warming (2, 3) and there are concerns that forests are reaching a tipping point beyond which they could switch irreversibly to open savanna systems (4). Forecasting the future of tropical forests is challenging, because little is known about the ways different species will respond to changing climate, or the resilience provided by that diversity (5–8). To understand the likely responses of forests to further climate change, ecosystem models need to represent growth and mortality processes of individual trees more accurately than is currently the case (9–11).

Remote sensing of individual upper-canopy trees can improve estimates of forest carbon (12) and provide a means of tracking growth and mortality over large spatial scales. Traditional monitoring approaches rely on measuring stem dimensions in permanent inventory plots, and periodically revisiting those plots to assess recruitment, growth and mortality (13). However, the coverage of such plots in the tropics is limited (~0.0002% of tropical forests are sampled by the main plot networks) and their locations are often dictated by ease of access rather than by robust statistical sampling designs (14–16). Furthermore, upper-canopy trees store the majority of carbon in tropical forests, but few of them are sampled in inventory plots (17–19). This under-sampling is particularly problematic when assessing impacts of climate change, because upper-canopy trees are most vulnerable to periods of water shortage (20, 21) which are increasing in frequency (22). Remote sensing has the potential to overcome these sampling challenges by providing wall-to-wall maps that can be used to monitor millions of upper-canopy trees.

Remote sensing of individual trees has mostly focused on airborne lidar data, which is used by the forestry industry to map trees at landscape scales (23). Delineating individual trees from airborne lidar datasets is most successful for conifer, because their apical dominance results in clear local height maxima that make tree crowns easily distinguishable (12, 24), but complex tropical canopies have presented a far greater challenge for lidar delineation (25). Tropical forest canopies are often densely packed with partially interwoven crowns which point-cloud clustering algorithms can struggle to distinguish (26). Furthermore, lidar surveys require expensive aircraft (airplanes, helicopters or high-end drones) and sensors whereas standard RGB imagery can be collected cheaply with drones.

Automatic delineation of trees in RGB photographs can harness colour and texture information to distinguish trees, even if they are structurally similar (27, 28). Most current methods of individual tree identification from RGB imagery use bounding boxes (Fig. S3) (29–31), but more exact delineation of the edges of tree crowns would provide information on crown area and lateral growth, and avoid mixing signals from neighbouring vegetation. Recent advances in neural network approaches to computer vision provide opportunities to recognise individual trees from standard digital photographs taken from drones. A class of machine learning algorithms called deep convolutional neural networks (CNNs) is revolutionising vegetation science through its ability to exploit spatial structures and automatically extract high-level features from image data (e.g. analyses of satellite imagery)(32–34). In the field of computer vision, exactly segmenting individual objects of interest from an image is known as *instance segmentation*. The Mask R-CNN algorithm has shown promise in tree crown identification and delineation in plantations (35, 36), pine forests (37, 38), urban woodlands (37) and forest fragments (39). Mask R-CNN has features that could allow it to overcome the challenges of delineating crowns in complex tropical canopies by discriminating based on the spectral and textural signals which are rich due to the phylogenetic diversity.

Here, we describe Detectree2, a system that automatically detects tree crowns from aerial RGB imagery. We adapted Facebook AI’s Mask R-CNN algorithm, (the *Detectron2* release), which has models that have been pre-trained on a wealth of available image data that can be transferred to new tasks (40, 41). We trained and evaluated Detectree2 on four tropical forest sites. In total, 3797 manually delineated tree crowns were used of which 1530 spatially separated crowns were reserved to evaluate the model. We evaluated the performance with F_1_ scores, which quantify the skill of the method in delineating individual tree crowns accurately. We expected a model trained at one site to drop in performance when transferred to making predictions of crowns at the other sites and that supplying a greater variety of training data would boost performance. As an example ecological application, we deployed the trained model across 14 km^2^ of airborne RGB imagery, automatically delineating 65,786 tree crowns. For context, this area is approximately 40% the total area of forest inventory plots in the main plot networks across the tropics (14, 15). We then combined these crowns with repeat airborne lidar data to investigate the growth and mortality rates of upper-canopy trees in relation to their height. Regional and global syntheses of forest inventory data suggest that growth slows down and mortality rates increase with tree size (42–46). We therefore expected to find the tallest trees we sampled to have lower growth rates and higher mortality rates than shorter trees. The Detectree2 Python package is available to install and apply on new regions^1^.

## Materials and Methods

### Study sites

The analyses were conducted at four locations across three tropical field sites:

1. Sepilok Forest Reserve (East and West), Sabah, Malaysia (5° 50’ N, 177° 56’ W)
2. Danum Valley Conservation Area, Sabah, Malaysia (4° 57’ N, 177° 41’ W)
3. Paracou Field Station, French Guiana (5° 16’ N 52 ° 55’ W)

Danum Valley hosts lowland tropical rain forests dominated by dipterocarp species that are among the tallest forests on the planet (47). The available data from Sepilok included ecologically distinctive areas to the East and West. Sepilok West consists mostly of tall forest (similar to Danum), while Sepilok East is a heath forest growing on shallow soils overlying sandstone, containing smaller, more densely packed trees (19). All three sites in Malaysia experience a similar climate with approximately 2300 mm rainfall per year with the wettest months between November and February (48). Paracou contains lowland tropical rain forest mostly on shallow ferralitic soils that lay on a variably transformed loamy saprolite (49). The mean annual rainfall is approximately 3000 mm with a three month dry season from mid-August to mid-November (50). See Sup. Note 1 for more details on the study sites.

### Remote sensing data

Airborne RGB surveys were conducted at all four sites using manned aircraft (Table 1). Repeat lidar surveys were also conducted at all four location (see Table 1, noting different sensors and altitudes between flights in Sabah). We analysed RGB imagery from 3.85 km^2^ of Malaysian forest, with a ground resolution of 10 cm. In Paracou, we sampled 10.2 km^2^ of imagery, with an 8 cm ground resolution. The raw imagery was orthorectified, georeferenced and collated into homogeneous mosaics using structure from motion in AgiSoft Metashape (51, 52) in Sabah. In Paracou the imagery was orthorectifed using TerraPhoto to the Canopy surface model derived from simultaneously acquired lidar data.

**Table 1.** Remote sensing data sources. The exact location of the sites is described in **Study Sites**. Resolution is given as ground resolution for the RGB imagery and as the processed CHM resolution for the lidar scans. Beam divergence is given at the 1/e^2^ points. Sepilok West and Sepilok East were separated for analysis due to the different characteristics of the Sepilok forest in these two areas.

| Scan dates | Region | Modality | Resolution | Pulse density | Beam divergence | Scanning angle | Altitude | Sensor |
| --- | --- | --- | --- | --- | --- | --- | --- | --- |
| 23-Oct-2014 | Danum | RGB | 10 cm | - | - | - | 796 m | Leica RCD105 |
| 01-Nov-2014 | Danum | Lidar | 1 m | 5 pls m<sup>-2</sup> | < 0.22 mrad | ± 14° | 2000 m | Leica ALS50-II |
| 19-Feb-2020 | Danum | Lidar | 1 m | 35 pls m<sup>-2</sup> | < 0.5 mrad | ± 30° | 200 m | RIEGL LMS-Q560 |
| 10-Oct-2014 | Sepilok | RGB | 10 cm | - | - | - | 796 m | Leica RCD105 |
| 05-Nov-2014 | Sepilok | Lidar | 1 m | 16 pls m<sup>-2</sup> | < 0.22 mrad | ± 14° | 2000 m | Leica ALS50-II |
| 15-Feb-2020 | Sepilok | Lidar | 1 m | 42 pls m<sup>-2</sup> | < 0.5 mrad | ± 30° | 200 m | RIEGL LMS-Q560 |
| 19-Sep-2016 | Paracou | RGB | 8 cm | - | - | - | 800 m | IXA180 Phase One |
| 19-Sep-2016 | Paracou | Lidar | 1 m | 35 pls m<sup>-2</sup> | < 0.25 mrad | ± 30 ° | 800 m | RIEGL LMS-Q780 |
| 15-Nov-2019 | Paracou | Lidar | 1 m | 35 pls m<sup>-2</sup> | < 0.25 mrad | ± 30 ° | 800 m | RIEGL LMS-Q780 |

### Manual tree crown data

To train and evaluate our automatic delineation approach, we created a manually labelled dataset of trees at all four sites. We generated our delineations using both RGB and lidar data and, in the case of Paracou, supplementary hyperspectral layers. We used several techniques to improve the accuracy of crown delineation, including manipulating the contrast and saturation of the RGB image to exaggerate differences between the crowns, and using a mask of the lidar data to remove irrelevant parts of the RGB imagery. These techniques meant that the vast majority of tree crowns were separable by eye but, it should be noted, that in rare cases, tree crowns were near impossible to delineate with certainty and the labeller’s best estimate was used. See Sup. Note 2 for further details.

We trained and tested our model with a total of 3797 manually delineated tree crowns across Paracou (1267), Danum (521), Sepilok West (1038) and Sepilok East (971). The crowns from Paracou were validated in the field with an expert local botanist, whereas the crowns in Malaysia were drawn by inspection of the remote sensing products. Four individuals performed the manual delineations which provided the network with variability in the inputs.

### Data preparation

The orthomosaics and corresponding crown polygons were tiled into squares of approximately 100 m x 100 m to be ingested into the network (40 m core area, 30 m overlapping buffers). To be included in the training and test sets, a minimum crown polygon area coverage of a tile was set at 40%. Including overly sparse tiles was likely to lead to poor algorithm sensitivity while being too strict with coverage would have limited the amount of training and testing data available.

If training and test crowns are close to one another, spatial autocorrelative effects are likely to inflate the reported performance (53). To avoid this, individual tiles (rather than individual crowns) were assigned to training and test sets ensuring spatial separation. Approximately 10% of the tiles from each site were reserved at random for testing. To avoid contamination of the test set, tiles with any overlap with the test tiles (including with the buffer) were excluded from the training set. The training tiles were further partitioned into 5-folds for cross validation. This allowed for the tuning of parameters and the implementation of early stopping (see **Training and model selection**) without exposing the test set. Details of the data processing are described in Sup. Note 3.

### Model architecture and parameterisation

Instance segmentation combines object detection with object segmentation. Once an object has been detected in a scene, a region of interest (as a bounding box) is established around the object. Then a “segmentation” is then carried out to identify which pixels within the region of interest make up the object of interest (and which lie outside; see Fig. S2 for an example).

We adapted Facebook AI’s Mask R-CNN architecture as it was the best in class algorithm upon release for instance segmentation when tested on the Microsoft COCO (Common Objects in Context) benchmark (40, 54) and has since been updated (as Detectron2) with improved training efficiency, documentation and transferability for integration into bespoke tools (41). We adapted the Detectron2 computer vision library to handle geospatial inputs/outputs and perform the delineation of individual tree crowns. The library performs instance segmentation by generating object “masks” which exactly circumscribe the objects in the image (see Fig. S2 for an example prediction). It also has a “model zoo"^2^ from which specific model architectures with weights from a variety of pre-training regimes can be loaded. Taking a pre-trained model (weights) and retraining it to perform a novel task (e.g. delineating trees from aerial imagery) is an example of *transfer learning* which can drastically reduce the amount of training data required to achieve acceptable performance on the new task (55). We selected the R101-FPN configuration^3^ as it has “the best speed/accuracy tradeoff” of the architectures available (41). Each object predicted by Detectron2 is associated with a confidence score which relates to how sure the network is in its prediction. A suitable threshold can be selected to optimise accuracy or balance precision and recall. Additional details are given in Sup. Note 4 and for full technical specifications, one should refer to the original papers and the Detectron2 repository (40, 41).

### Training and model selection

Training, tuning and model selection were performed with the five folds of training data tiles (see **Data preparation**). To test the effect of volume and diversity of training data on performance we employed three training regimes: (1) Training on data of a single site, (2) Training on all sites (“combined”), (3) Training on all sites and then trained with a fixed training period on the single site. The idea behind (3) was to train on the full available data and then ‘hone’ the delineator based on the local context.

Typically, a deep CNN would require several thousand training examples to learn a new task. This is a challenge in the case of tree crown delineation as manual delineation is time consuming. The burden was reduced by transferring a model trained on a different instance segmentation task (54). Additionally, the training data were augmented by applying several randomly applied transformations to the training cases including vertical and horizontal flips, rotation, and varying the saturation and contrast of the image.

The model hyperparameters (see Table S1) were tuned with a Bayesian hyperparameter sweep implemented on wandb.ai^4^. This is an automated process that allows an automated agent to iteratively adjust hyperparameters to optimise accuracy. The best performing models and optimal confidence threshold for a given model (see **Model architecture and parameterisation**) were selected based on the F_1_ score (see **Performance evaluation**) on the validation fold.

See Sup. Note 4 for more details on model architecture, training and validation. The Colab (Jupyter) notebooks in the GitHub repository^5^ illustrate the best practices for training and selecting models.

### Performance evaluation

After tuning and training, the best performing models were taken forward to be evaluated against the test tiles. Matches between predictions and manual crowns (i.e. true positives) were identified by assessing the degree of spatial overlap between possible pairs. A minimum area threshold for valid crowns was set to 16 m^2^. This removed fewer than 2% of manual crowns and introduced a level of consistency between sites and between the effort given by the manual delineators. The threshold was small enough to allow for an inclusive analysis of the variation in performance by tree height. Crown overlap was calculated as the area intersection over union (see Fig. 1):

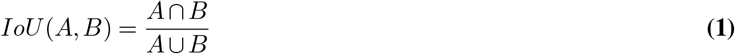

where *A* is the automatically delineated crown area and *B* is the manually delineated crown area. An *IoU* of an overlapping pair of more than 0.5 was considered a match. This is a commonly used threshold in similar studies (e.g. (25, 36)) that allows for small discrepancies in alignment and outline. These “true positives” as well as the unmatched predictions (false positives) and unmatched manual crowns (false negatives) were used to calculate the precision, recall and F_1_ score of the automatic predictions.

**Fig. 1.**
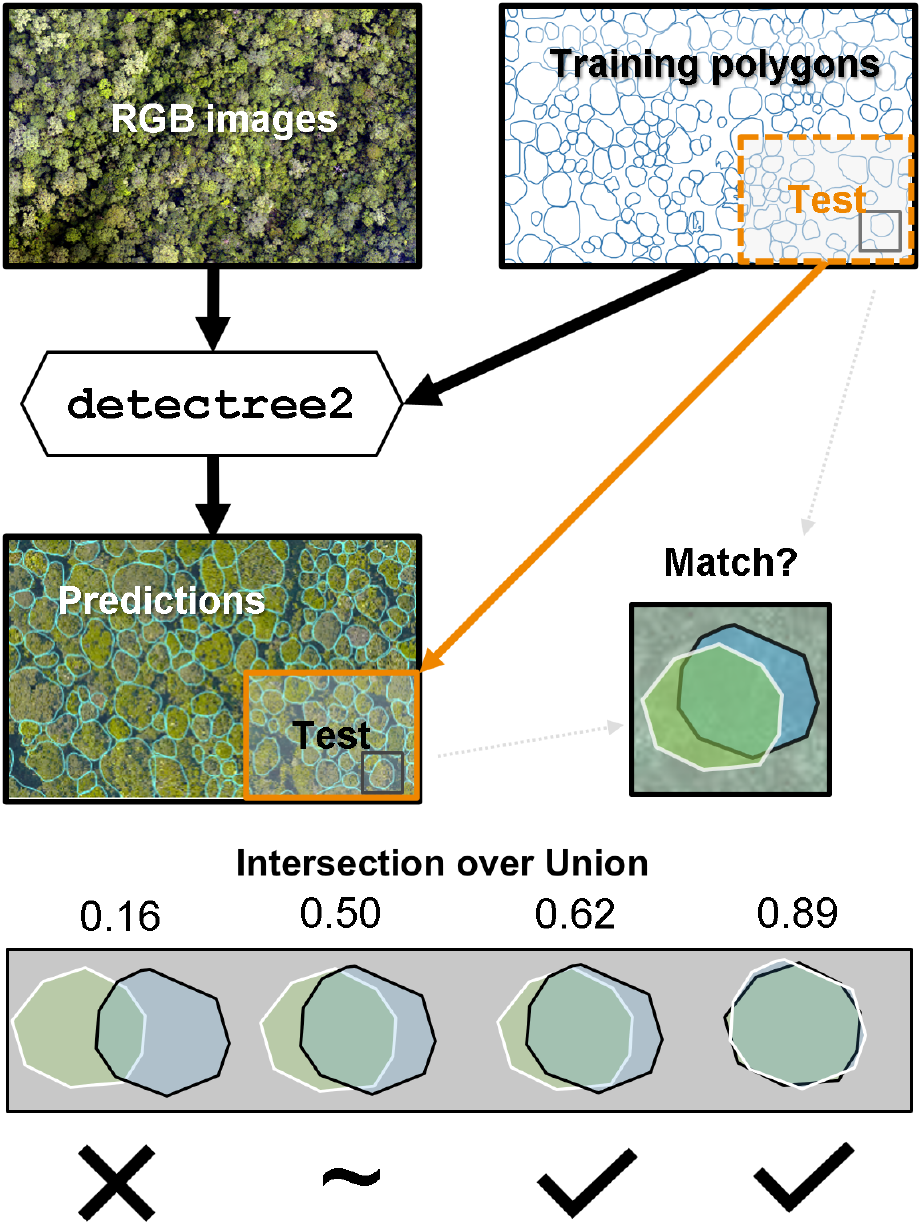
The automatic tree crown delineation workflow. Manually delineated crowns are randomly split into training and test sets (though the figure suggests that the sets were determined geographically, this is purely for visual clarity). The Mask R-CNN framework combines the training set with the RGB imagery to learn how to delineate automatically from RGB images. A set of automatic predictions are produced across the entire RGB image and compared to the test set to evaluate the performance of the automatic delineations. Intersection over union (*IoU*) is used to determine when an automatic crown has been successfully matched with a manual crown.

Despite the best efforts of the manual delineators and selecting for tiles with high manual crown coverage, the manual crowns were inevitably an incomplete representation, so recall (fraction of relevant instances retrieved) was an insightful metric. However, to ensure balance with precision we used the balanced F_1_ score metric to assess and compare the accuracy of the models. This approach is not biased by tree crown area and is widely used in tree crown segmentation studies (36, 39). See Sup. Note 5 for more details on the evaluation metrics.

To evaluate the performance of Detectree2 across tree heights, we assigned a height to each test crown (based on the median pixel value of the initial CHM within the crown) and arranged them into 5 m height bins. The shortest bin (0-5 m) at each site was iteratively merged with the next shortest bins until more than 10 individuals were represented (e.g. at Paracou 17 trees with a median height of 22.97 m fell into the lowest bin of 0-25 m). An equivalent process was used to define the highest bin at each site. The median tree height was calculated within each bin.

### Transferability across sites

To determine whether models were able to generalise across different tropical forest areas, we evaluated the performance of the models when trained on one site and transferred to others. We compared these performances against the effect of using the “combined” training regimes described in **Training and model selection**.

### Application to monitoring growth and mortality

We applied our best models for each site to their entire tiled orthomosaics (excluding the very edges where distortion is prominent) to generate site wide crown maps. We combined these crown delineations with repeat lidar surveys to determine the height changes in individual trees in our four sites. We determined the relationship between tree height and tree growth by fitting a robust least squares regression (56) to the data. Robust least squares was chosen to minimise the effects of outliers and mortality events on the regression. We note that here we are measuring the vertical growth of trees, instead of the growth in diameter at breast height (DBH) which is traditionally measured in forest inventory data.

To estimate mortality rates, we needed a suitable metric to identify mortality events. We took a statistical approach defining a mortality event as a negative change in height of more than three standard deviations below the robust least squares fit. This allowed for the possibility that a mortality event may uncover another layer of vegetation rather than the forest floor. This choice was ratified by manual inspection of trees meeting this threshold, and confirming that they constituted mortality events. Annual rates were determined by dividing by the time between lidar scans.

Differences in lidar scanning parameters (pulse density, scanning angle, flight height etc.) can bias height estimates (57). For this reason, we resisted reporting a direct comparison of reported growth and mortality rates between sites. As our focus here was on demonstrating the use of Detectree2 for locating crowns, we considered that a detailed exploration of the potential biases from the lidar data beyond the scope of the current paper.

### Computation

Training deep CNN models can be computationally expensive and benefits from the availability of GPUs. Model training and evaluation was performed on the Google Colab (Pro) platform which employed Intel(R) Xeon(R) CPU at 2.30GHz with 12.8 GB RAM and Tesla P100-PCIE-16GB GPUs. On this platform, model training always completed within 2 hours.

## Results

### Performance by site and tree height

Detectree2 located and delineated trees well (F_1_ > 0.56) across all sites (see Table 2). It performed better in the tall dipterocarp dominated forests of Danum and Sepilok West and worse in the more compact forests of Sepilok East and Paracou. Indeed, Danum, the site with the best performance, had the greatest proportion of the tallest class of trees of any of the sites (see **Extended Results** for a full table of results). There was no apparent relationship between the amount of training data available at a site and the performance of the automatic delineator suggesting forest structure was the key determinant of accuracy. Where predictions were not accurate, it was slightly more likely to be from under-segmention (0.23-0.45) than over-segmentation (0.13-0.23) (58) across all sites (see **Extended Results**).

**Table 2.** Precision, recall and F_1_ score of Detectree2 tree crown delineations by site as measured against the manual crowns of the test set tiles. The unweighted means of the metrics across individual sites are given as a summary overall performance.

| | # test trees | Precision | Recall | $F_1$ score |
| --- | --- | --- | --- | --- |
| Paracou | 381 | 0.595 | 0.543 | 0.568 |
| Danum | 278 | 0.713 | 0.662 | 0.687 |
| Sepilok East | 167 | 0.612 | 0.653 | 0.632 |
| Sepilok West | 704 | 0.640 | 0.656 | 0.648 |
| *Average (sum)* | *(1530)* | *0.640* | *0.629* | *0.634* |

Across all sites, accuracy improved with tree height (see Fig. 3). This is likely due to the increased crown visibility of tall trees in the RGB images. Paracou has the least well differentiated canopy of all sites which may explain relatively poor performance there.

**Fig. 2.**
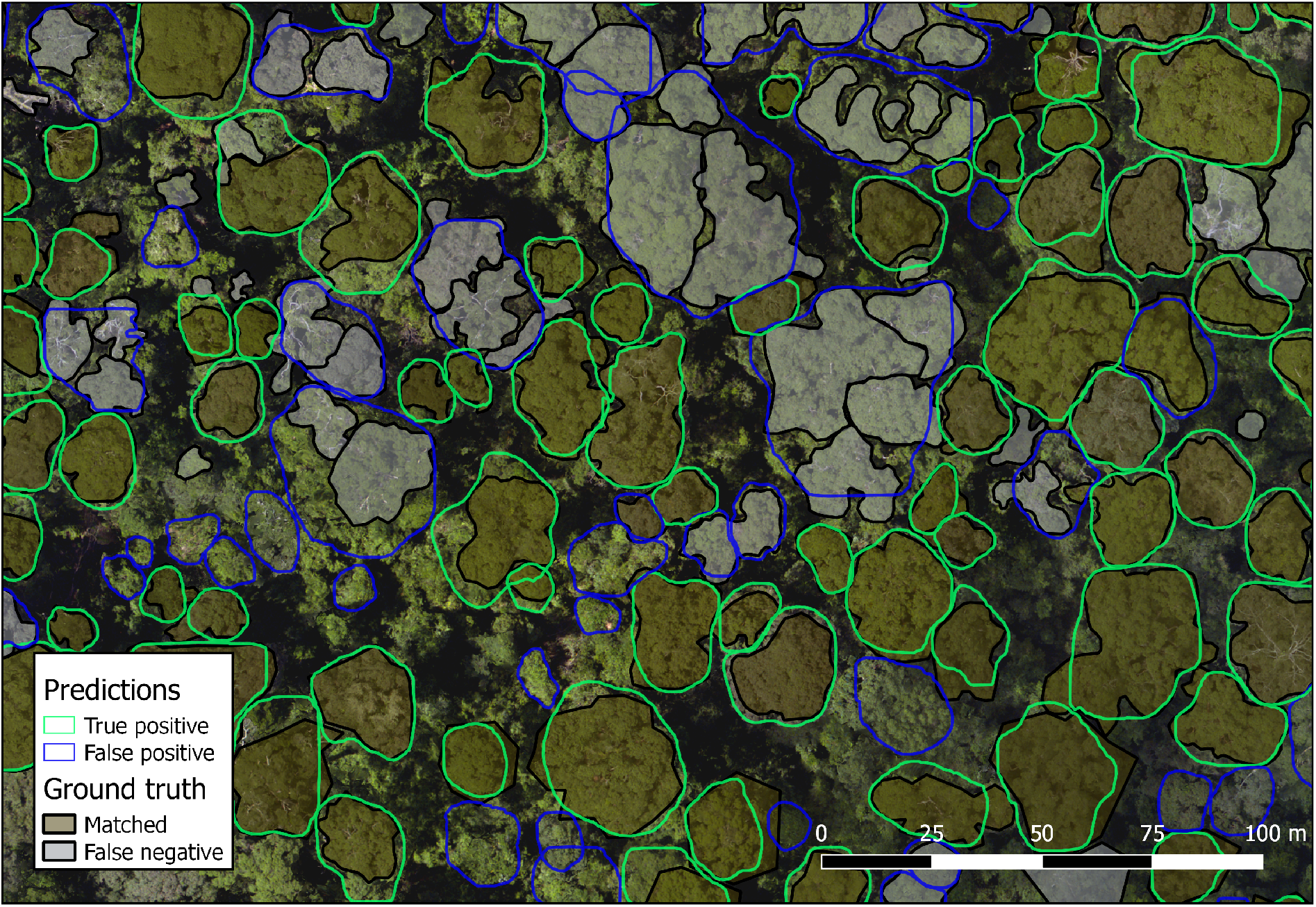
An area of predicted crowns (transparent) overlaid on ground truth crowns (shaded with black outlines) at Danum. Colors and shading are used to indicate whether individual crowns have been successfully delineated. Some examples of under-segmentation (where a single prediction encompasses multiple ground truth crowns) and over-segmentation (where multiple predictions false try to split a single ground truth crown) are visible.

**Fig. 3.**
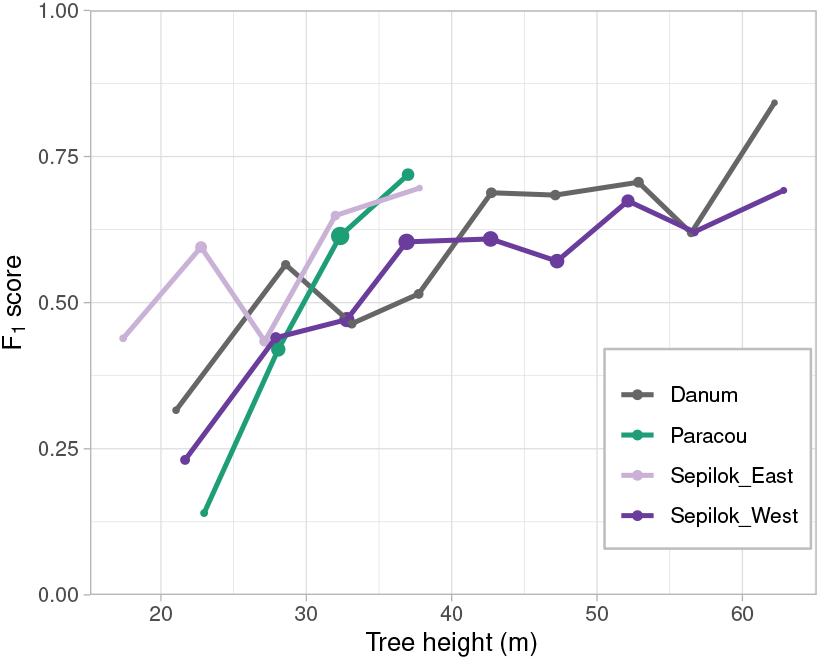
F_1_ scores of the tree crown delineations at the four different sites across tree heights. Bins of 5 m width were used to calculate F_1_ score and corresponding median tree height. Point area is scaled by the number of test trees in the bin.

### Performance between forest types

Danum and Sepilok West have tall dipterocarp dominated forests whereas Paracou and Sepilok East have a more compact forest structure. As we expected, performance degrades when testing a model on a different forest type to the one it was trained on (see Fig. 4a). For example, the performance at the forests of Sepilok West is significantly degraded when a model trained on Sepilok East or Paracou is used. In contrast, there is no drop in performance for predictions at Danum when the Sepilok West model is used and there is even a slight increase in performance for Sepilok East predictions when the Paracou model is used.

**Fig. 4.**
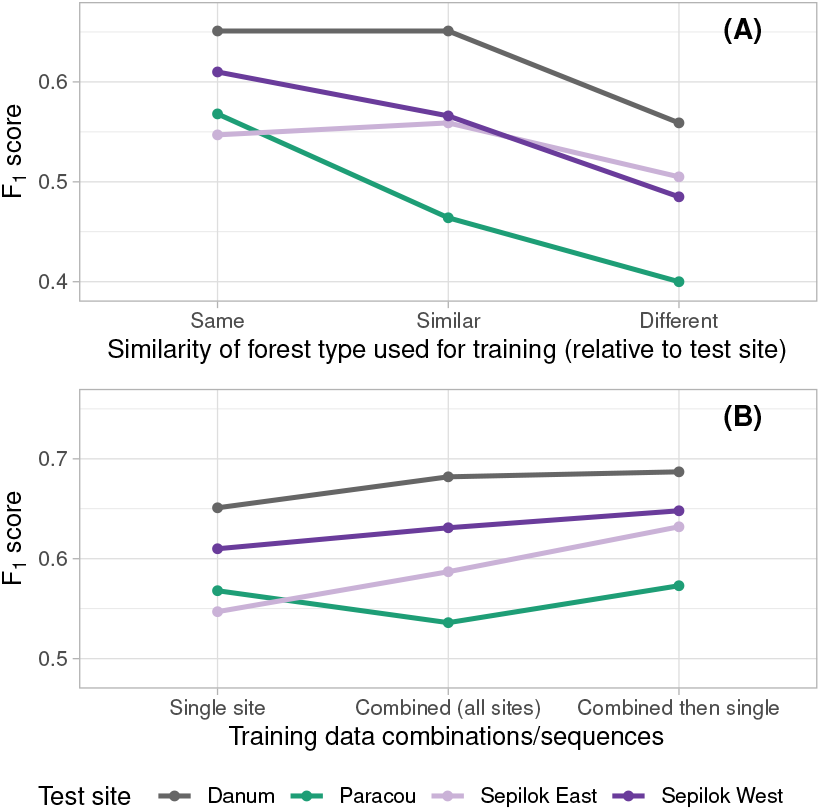
Sensitivity of Detectree2 delineation accuracy to the training data used. In **(A)**, “Same” indicates that training and testing took place at the same site. Sepilok West and Danum are “similar” forest types in that they are tall dipterocarp dominated forests in contrast to Sepilok East and Paracou that are shorter forests with a larger number of trees per hectare. As each site has two “different” sites and an average was calculated for the F_1_ score. **(B)** shows the change in performance that occurs through employing different combinations of the training data. That can be just a single site, all sites at once (“combined”) or all sites at once followed by a limited number of iterations on the site to be tested on.

In general, the model that was trained on all sites at once (“combined”) outperformed the models that were trained on just a single site with the exception of Paracou (see Fig. 4b). Across the board, the best performing models were those that were exposed to data from all sites before being trained for a fixed number of iterations at the site to be predicted on. This suggests that providing a broad range of input data helps the networks to learn the key visual features but further tuning for local context helps maximise performance.

### Application: Growth and mortality

One application of Detectree2 is to study tall tree growth and mortality rates. To do this, we overlaid Detectree2’s tree crown predictions at the start date for each site on repeat lidar data (as canopy height models described in Sup. Note 1) to retrieve the tree height dynamics over time.

We were able to estimate the relationships between tree height and tree growth for each site by fitting robust least squares linear relationships between the two variables for Danum, Paracou, Sepilok East and Sepilok West Fig. 5. The regression coefficients and intercepts are given in Table S2. The growth rate decreased with tree height in all sites.

**Fig. 5.**
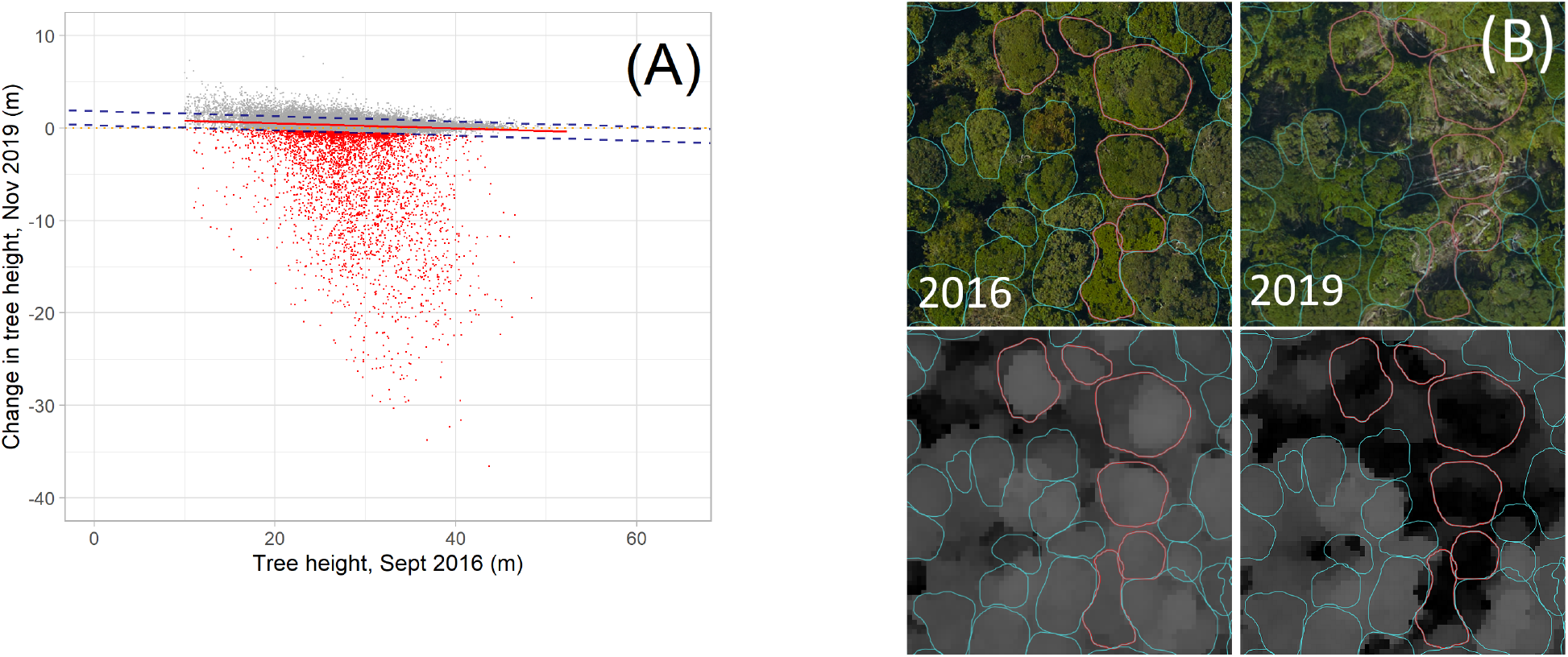
**(A)** shows the robust least squares fit for change in height and tree height for Paracou. The dashed lines indicate three standard deviations either side of the best fit and red points below the lower bound indicate likely mortality events. **(B)** illustrates how predictions were overlaid on lidar data, and shows mortality events clearly visible in the lidar and the RGB imagery. The 2016 imagery is shown on the left, 2019 on the right. Crown delineations are based on the earlier imagery.

We assumed trees had died when their height decreased substantially. To evaluate this quantitatively, we fitted a robust least squares to the height change, against the original height of the tree, taking trees that were 3 standard deviations below the mean of the regression fit to be mortality events. The robust least squares regression differs from ordinary least squares as outliers contribute less to the regression fit. Therefore the robust least squares weights the fit towards those trees which did not suffer large height loss, and by taking the threshold to be 3 standard deviations we aim to identify only those trees that are outside the assumed normal distribution of typical tree growth and measurement error. Furthermore, as the robust least squares still incorporates outliers when fitting the data, three standard deviations was considered sufficient to identify mortality events. Fig. 5 illustrates how certain trees were identified as mortality events and some visual examples of mortality events from Paracou are given alongside (see Fig. S12 for the other sites).

The mortality rates increased with tree height Fig. 6. The given uncertainty estimates were determined by bootstrapping. A detailed table of the growth and mortality rates can be found in **Extended results**.

**Fig. 6.**
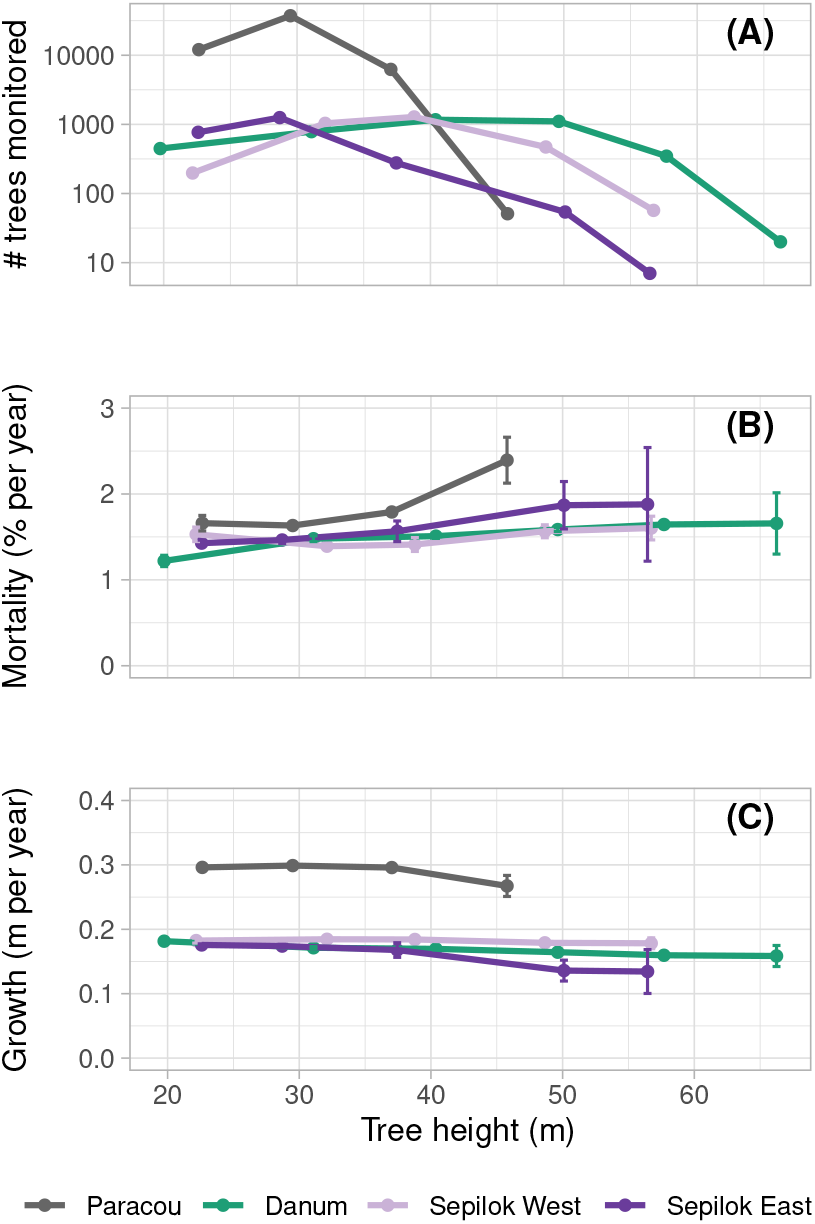
**(A)** shows the distribution of tree heights per site. **(B)** shows the mortality rates of trees of different heights in each site, and **(C)** gives the growth rate of trees split by height bin. Due to biases in tree height measurements that can arise from differences in lidar scan parameters we advise against a direct comparison of growth and mortality rates between sites. Uncertainty estimates were determined by bootstrapping.

## Discussion

### Improved tropical crown delineation

Accurately delineating trees in remote sensing data is a long-standing problem in ecology and conservation, and would enable us to efficiently monitor large areas of forests. Detectree2 addresses this problem, delineating individual trees in aerial RGB imagery with high precision and recall. We used Detectree2 to automatically delineate 65,786 trees across three tropical forests. We found that the accuracy of Detectree2 increased with tree height, meaning that that the tall trees which store the most carbon are also the most reliably delineated.

Detectree2 performed well across a range of challenging, dense, closed canopy forests. It is able to exactly delineate highly irregular crowns within the jigsaw of the canopy rather than simply identifying a bounding box. This opens up new opportunities for tracking dynamic processes including growth and demographics (as demonstrated here) as well as phenology (where bounding boxes would risk mixing signals). Furthermore, Detectree2 is relatively accessible since it requires a low number of manually delineated trees as training data compared to other methods (30, 39). These advantages are partly due to Detectree2 being built on a state-of-the-art pre-trained model. While direct comparison is impossible due to the different test data and the differing tasks (instance detection vs. segmentation), our method performs comparably to the results of (30), which reported a tree crown recall of 0.69, a precision of 0.61 and an F_1_ score of 0.65. Our results did not match the Mask R-CNN performance reported in (39) but this study is based on semi-synthetic images (i.e. constructed by stitching together existing images) of forests and so is not directly comparable.

### Generalisability across sites

There was no obvious relationship between the amount of training data available at a site and the accuracy attained there. Rather, forest type and tree height distribution seemed to be the key factors for determining accuracy. The well differentiated forest at Sepilok West and Danum were the easiest to delineate while the lowest accuracy was in Paracou which has little variation in the height of the visible canopy. Furthermore, at Paracou it is common to observe crowns mixing and growing into each other which makes visually separating the crowns challenging. This in turn is down to soil type and other biogeographic factors.

We found that the accuracy dropped when transferring a model trained on one forest type to predict on another. However, we found that Detectree2 can be quickly trained to perform well on new areas of forest using around 10 images (each ~1 ha in scale) with all visible tree crowns manually delineated. This manual delineation represents approximately 4 hours work. The best performing models were those that were exposed to training data from all the sites and then “honed” with a limited number of training iterations on the site to be predicted on. This suggests that our trained models (provided freely with the Python package^6^) could be transferred to a new site with very little manual data or training iterations.

We note that the manual delineations were done by different people focusing on different parts of the sites. There was no clear effect of different delineators on the results but this would be somewhat confounded with site differences.

### Application: Growth and mortality rates

Tall trees store the majority of forest carbon and dominate many important forest nutrient cycles. However, they are rare and therefore poorly represented in traditional field inventories (46) which makes estimating their growth and mortality rates particularly challenging (42–45). Tall trees are also particularly sensitive to the effects of climate change, such as increased wind speeds and drought (21), and as such tracking their dynamics over time is increasingly important. Recent remote sensing studies are bringing new insights into disturbance patterns by mapping the gaps left in the forest canopy after a tree (or multiple trees or branches) have fallen (59–61). Tracking individual trees over time instead of gaps will make it easier to interpret our results in an ecological context and also to compare the results more directly to the available field inventory data.

Across all sites taller trees had higher mortality rates and lower growth rates. This aligns with large scale analyses of field-based studies (42). The apparent higher growth and mortality rates in French Guiana as compared to the sites in Malaysia was potentially a result of biases introduced to the variation in scan parameters (flight height, pulse density, time of year etc.) and so the values should not be directly compared across sites. Inventory data shows that mean DBH growth for trees at Paracou was 1.2 mm/yr (62) compared to 0.9 mm/yr in Sepilok East, 1.1 mm/yr in Sepilok West and 0.5 mm/yr in Danum (10, 63). We note that these inventory measured DBH growth rates may not be directly comparable to the height growth measured in this study. Another caveat is that we defined mortality as a drop in height of more than a statistically determined threshold. We do not verify directly that the tree has died, although it is likely that it has snapped or uprooted. Further analysis would help to understand the discrepancy in observed height change at Paracou in comparison to the other sites but is not the focus of the current study. Nevertheless, we believe this example application demonstrates the utility of Detectree2 in expanding the sample of trees under observation.

### Future methodological developments and applications

Detectree2 performs impressively when delineating tall trees but it fails to delineate a significant proportion of trees. There is considerable scope to increase the quantity and variety of training data by labelling more trees by hand. A more robust approach to compensating for shadowed regions may also support the detection of trees otherwise obscured by their neighbours.

The fact that Detectree2 can be quickly trained to perform well on a new type of forest and imagery demonstrates that it is a useful tool for forest management. Many conservation or restoration projects have access to low-cost imagery from drones or satellites. Detectree2 would allow them to quickly quantify and track the number and size distribution of trees across an entire landscape. In combination with other remote sensing data sources, this could allow for improved carbon stock and dynamics estimation. Estimating carbon stocks in forests has traditionally been done using area-based methods which discard considerable granular information at the individual tree level (19).

We focused on aerial RGB imagery which is the cheapest and most widely available imaging source for tropical forests. We also benefited from the variety of pre-trained models that come with this data type. However, different data sources may provide additional information that would help to discern differences between crowns. In particular, multi-spectral imagery that typically includes additional bands in the near-infrared is commonly used to study differences between trees due to the optical properties of vegetation (64). Alternatively, the canopy surface (a raster expressing the height of the canopy) is commonly used in traditional segmentation techniques (e.g. watershed algorithms) and is produced with photogrammetry as a step in generating an orthomosaic. Including this as a layer would add an additional dimension of information that could help to distinguish fine differences in structure. It would be straightforward to include additional (or different) bands to the Detectree2 framework but it would forego the utility of the pre-trained models. Therefore, it is likely that significantly more training data and computational resources would be required to train a model (from scratch) to the desired performance.

Ideally, we could apply this approach to satellite imagery to perform global analyses. Preliminary tests suggest that Detectree2 can accurately delineate trees in RGB imagery at 2 m resolution (see Sup. Note 7) which is equivalent to modern high-resolution satellite imagery. If this proves possible it will help answer many long-standing questions in forest ecology as well as provide an important tool for forest management. A “random resizing” augmentation step would further help improve generalisability across resolutions and incorporating “small object” detection features (65) would improve the sensitivity to shorter trees.

While we studied the delineation of a single class (tree), Detectree2 can be trained and make predictions on multiple classes. This may allow for low-cost species identification and mapping. It may also help to automatically assess liana infestation occurrence. Previously, hyperspectral data has been employed to address this problem but with limited success due to the phylogenetic and spectral diversity of lianas and relatively low spatial resolution of hyperspectral imagery (66, 67). The availablility of Detectree2 as an open-source Python package means other research groups can test its efficacy on their own research questions.

## Supporting information

Supplemental results

## ACKNOWLEDGEMENTS

We thank the NERC Earth Observation Data Acquisition and Analysis Service (NEODAAS) for supplying data and computational resources for this study. We thank CEDA for the 2014 Sepilok and Danum remote sensing data. SHMH received funding from the Centre for Doctoral Training in Application of Artificial Intelligence to the study of Environmental Risks (AI4ER, EP/S022961/1), which is supported by the Engineering and Physical Sciences Research Council (EPSRC). JGCB was supported by the NERC C-CLEAR doctoral training programme [PDAG/501]. TDJ and DAC were supported by NERC grant [NE/S010750/1]. DAC was supported by the Franklinia Foundation. We thank Stephen Goult for contributing to discussions. Data collection in French Guiana was supported by CNES who funded the 2016 hyperspectral, RGB and lidar data over Paracou and Labex CEBA (ANR-10-LABX-25) for contributing financial resource for the field validation of manual crown segmentations. The 2019 data in Paracou and 2020 data in Sabah were funded by NERC [NE/S010750/1]. The 2014 Sabah data were also funded by NERC [NE/K016377/1].

## AUTHOR CONTRIBUTIONS

SHMH, JGCB, TDJ and DAC conceived the idea and designed the study. JGCB led the development of the Python package with contributions from JH, MA and SHMH. WJ processed the Malaysian aerial photographs. MA-K, GV and JGCB conducted the Paracou data collection and processing. GV, XJK, SHMH, JGCB and TDJ manually labelled tree crowns and conducted the analysis. SHMH, JGCB and TDJ wrote the manuscript and all authors contributed to the final version.

## DATA AVAILABILITY

Data will be made available on Zenodo upon acceptance.

Airborne LiDAR and RGB imagery collected in Malaysia in 2020 are available here: http://dx.doi.org/10.5285/dd4d20c8626f4b9d99bc14358b1b50fe

## Word Counts

This section is *not* included in the word count.

### Notes on Remote Sensing for Ecology and Conservation

- Maximum of 5000 words, excluding acknowledgements, references and figure and table legends.
- Title page, including a concise and informative title, authors’ names, authors’ affiliations, and contact information
- Running title not exceeding 45 characters
- Word count of the entire paper broken down into summary, main text, acknowledgements, references, tables and figure legends
- Number of tables and figures
- Abstract (maximum 300 words) and 4–6 keywords
- Cover letter detailing the key findings, the novelty of the work and how the manuscript fits the aims and scope of the journal
- Text (Introduction, Materials and Methods, Results, Discussion)
- Acknowledgements, including details of funding bodies with grant numbers
- Data accessibility
- Literature cited (see below for tips on references)
- Figure legends
- Tables (may be sent as a separate file if necessary)
- Figures
- The manuscript should be double spaced and line numbered throughout

### Statistics on word count

~~~
File: Article.tex
Encoding: utf8
Sum count: 6043
Words in text: 5366
Words in headers: 85
Words outside text (captions, etc.): 587
Number of headers: 23
Number of floats/tables/figures: 8
Number of math inlines: 4
Number of math displayed: 1
Subcounts:
  text+headers+captions (#headers/#floats/#inlines/#displayed)
  126+16+0 (1/0/0/0) _top_
  298+1+0 (1/0/0/0) Abstract
  842+1+0 (1/0/0/0) Section: Introduction
  0+3+0 (1/0/0/0) Section: Materials and Methods
  185+2+0 (1/0/0/0) Subsection: Study sites
  108+3+67 (1/1/0/0) Subsection: Remote sensing data
  199+4+99 (1/1/1/0) Subsection: Manual tree crown data
  205+2+0 (1/0/0/0) Subsection: Data preparation
  302+4+0 (1/0/0/0) Subsection: Model architecture and parameterisation
  264+4+0 (1/0/0/0) Subsection: Training and model selection
  379+2+0 (1/0/3/1) Subsection: Performance evaluation
  48+3+0 (1/0/0/0) Subsection: Transferability across sites
  274+6+0 (1/0/0/0) Subsection: Application to monitoring growth and mortality
  54+1+0 (1/0/0/0) Subsection: Computation
  0+1+0 (1/0/0/0) Section: Results
  174+6+146 (1/3/0/0) Subsection: Performance by site and tree height
  201+4+123 (1/1/0/0) Subsection: Performance between forest types
  319+4+151 (1/2/0/0) Subsection: Application: Growth and mortality
  0+1+0 (1/0/0/0) Section: Discussion
  260+4+0 (1/0/0/0) Subsection: Improved tropical crown delineation
  260+3+1 (1/0/0/0) Subsection: Generalisability across sites
  346+5+0 (1/0/0/0) Subsection: Application: Growth and mortality rates
  522+5+0 (1/0/0/0) Subsection: Future methodological developments and applications
~~~

## Supplementary Note 1: Study sites and remote sensing data collection

Orthomosaics were generated from the raw aerial photographs in AgiSoft Metashape. This software uses structure from motion to generate a 3D elevation surface from the raw aerial photographs (51) which is then projected back into 2D to create a landscape scale image (orthomosaic). This gave georeferenced photographs, largely corrected for distortions. Some distortion remained at the edges of the mosaic, so the analysis focused on the core areas where the imagery was not distorted.

### A. Sepilok (East & West) and Danum

Sepilok Forest Reserve (5° 50’ N 117° 56 ‘ E) is a region of lowland tropical rainforest in Sabah, a state in Malaysia. The Reserve is one of the oldest protected areas of forest in Asia, founded by the Sabah Forest Department in 1931. The Reserve spans nearly 4500 ha, with ground elevation varying between 50 and 250 metres above sea level. Three distinct forest types are present: alluvial dipterocarp, sandstone dipterocarp and heath forest. Danum Valley contains lowland tropical rain forests dominated by dipterocarps and are among the tallest forests on the planet (47). The three Malaysian sites experience a similar climate with approximately 2300 mm rainfall per year with the wettest months between November and February (48).

The 2014 lidar data of Sepilok and Danum were collected using a Leica ALS50-II ALS flown at an altitude of nearly 2000 metres, attached to the belly of a Dornier 228-201. The ALS sensor emitted pulses at around 80 Hz with a field of view of 14.0°, and a footprint of about 40 cm diameter. The average pulse density was 11 pulses m^-2^. The ALS data were preprocessed by NERC’s Data Analysis Node and delivered in LAS format.

The 2020 lidar data in Sepilok was collected using a RIEGL LMS-Q560 mounted on a helicopter flying at 200 metres altitude at a ground speed of approximately 100 km/hr. This resulted in an average pulse density of 38 m^-2^. Further processing used LAStools (http://rapidlasso.com/lastools/). Points were randomly resampled to match the point density across dates. Points were split into two groups, ground and non-ground, and a digital elevation model (DEM) was fitted to the ground points, producing a raster of 1 m resolution. The DEM elevations were subtracted from elevations of all non-ground returns, known as the digital surface model (DSM) to create a canopy height model (CHM) raster of resolution 1 m.

The 2014 RGB imagery was collected using a Leica RCD105 Digital Camera, attached to a plane flown at an altitude of 796 metres, with a ground resolution of 10 cm. Photographs have been orthorectified and collated into homogenous mosaics using Agisoft Metashape. This software used the Structure from Motion method to calculate the elevation of the observed surface, of which the photographs are mapped to, This results in georeferenced photographs that are corrected for distortions. The files were delivered in the GeoTIFF format.

### B. Paracou

The Paracou field station is situated in a lowland tropical forest near Sinnamary, French Guiana (5°16’N 52 °55’W). It is a forest similar to Sepilok Forest Reserve, which is also a lowland tropical forest.

ALS data were acquired in September 2016, and November 2019 by ALTOA, operating a RIEGL LMS-Q780 sensor attached to an aircraft flying at 800m. On all dates, the scan frequency was 400 kHz and the final point density was above 50 m^-2^. The creation of a CHM was carried out using the same process as used for Sepilok.

On the same flights as the ALS scans, RGB images were collected using an IXA180 Phase One camera with an 8 cm ground sampling distance.

Hyperspectral imagery was also collected over Paracou, with a Hyspex VNIR-1600 (Hyspex NEO, Skedsmokorset, Norway) sensor-mounted alongside the RIEGL scanner. Its bands covered a spectral range of 414–994 nm with a spectral sampling distance of 3.64 nm. Images were orthorectified and georeferenced to 1 m spatial resolution with the PARGE software using the DSM from the lidar data.

## Supplementary Note 2: Tree crown data

### A. Manual delineation

Manual tree crown labelling was carried out at each study site, to create training and test sets for our analysis. Labelling was carried out in open-source graphical geospatial software, QGIS (68), using a combination of RGB, lidar, and, in the case of Paracou, hyperspectral imagery. To increase the contrast between individual trees the colours of the RGB imagery were stretched. Polygons were carefully drawn around the perimeter of all distinguishable trees for a number of areas in each region. JGCB carried out the manual labelling for Paracou. A visualisation of the difference between the original and the stretched image is given in Fig. S1.

### B. Training and validation data

Examples of the training data are given in Fig. S4.

## Supplementary Note 3: Data preparation and processing

To apply Detectree2 to remote sensing RGB images of tropical forests, the geospatial raster image must be tiled and converted into a png format, and the manual crown segmentations used for training must also be converted from geospatial shapefiles into JSON files that align with the pngs. These training tiles are ingested into Detectree2, which then learns how to identify a tree, and then can make predictions on new tiles.

**Fig. S1.**
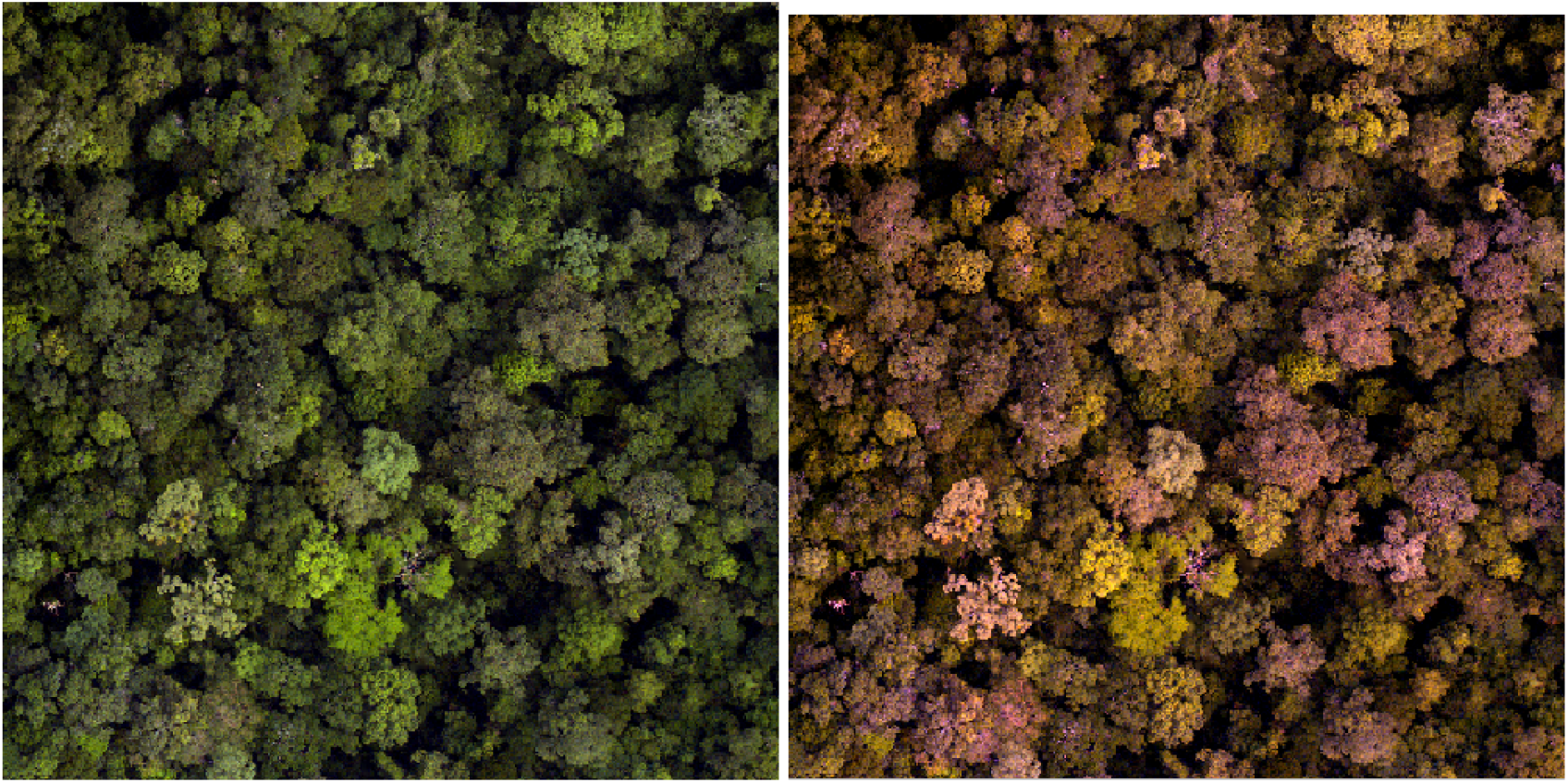
An original RGB image is displayed on the left, while a stretched RGB image is displayed on the right of this figure. Stretching the colours of the image to the values allows for easier identification of individual trees when carrying out manual tree crown delineations. The effect is particularly noticeable in the lower left corner of the images.

Training tiles were separated into 5-folds so that cross validation could be performed during the hyperparameter tuning phase. Testing tiles were spatially separated from the training tiles and only seen by the network at the evaluation phase.

## Supplementary Note 4: Model architecture, tuning and training

### A. Model architecture

Mask R-CNN (40) is a framework for instance segmentation. It builds on Faster R-CNN (69), which carries out object detection using a Region Proposal Network (RPN). An RPN is a fully convolutional network, trained end-to-end, that generates Regions of Interest (RoIs) for each image. These RoIs have object bounds and objectness scores attached to them, giving the bounds of the RoI, and the likelihood of the RoI containing an object. These RoIs are then passed through fully connected convolutional networks to determine the class of image contained, and the exact mask of each object, within the respective bounding box. For full details on the structure of Mask R-CNN, please refer to the GitHub repository for this work (https://github.com/PatBall1/Detectree2). The schematic of the architecture of the model architecture is given in Fig. S2.

We selected the R101-FPN configuration - this architecture combines a 101 layer deep ResNet (70) module with a Feature Pyramid Network module. The configuration sets the *backbone* of the network which is the part that views and extracts features from the scene as a whole. The R101-FPN backbone consists of a 101 layer deep ResNet (71) module with a Feature Pyramid Network (72) module. The initial model weights were generated from pre-training of the network on the ImageNet dataset^7^. It is possible to “freeze” the backbone to different depths depending on the amount of flexibility the user wants to introduce in moving away from the pre-trained model weights. An example of the different predictions is given in Fig. S3.

### B. Data augmentation

The training data was augmented by submitting the training data to a variety of transformations including vertical and horizontal flips, rotation, and varying the saturation and contrast of the images. These augmentations are designed to increase the variety of training data seen by the model, to allow it to generalise more readily to new images.

### C. Training and hyperparameter tuning

based on a given accuracy metric. We selected the *AP50* of the segmentation predictions on the (randomly selected) validation fold as our metric for optimisation. AP50 is the average precision of predictions when a correct match is granted for *IoU* > 0.5.

AP50 was also used as the metric for *early stopping* whereby if the model performance failed to improve for a set number of training iterations (the “patience”), training would be terminated and the best model up to that point would be saved. This is a technique for regularisation and prevents wasted computation. As with all deep networks, there were several of hyperparameters that controlled the way the networks handled data handling and were trained. The first of these relates to the architecture of the algorithm, namely the number of hidden layers in the ResNet backbone (71) of the network. ResNets are used as the backbone of Mask R-CNN as very deep neural networks are particularly difficult to train, due to the vanishing gradients problem (73). ResNets use skip connections, whereby certain layers in the network are skipped during backpropagation to avoid this problem.

Other hyperparameters that can be optimised in Mask R-CNN relate to the training of the network. Namely the learning rate, the number of iterations and the batch size. The optimisation of these hyperparameters was carried out using Weights and Biases, which uses grid search to determine the optimal hyperparameters. The hyperparameters selected are given in Table S4.1.

**Table S1.** Tunable hyperparameters (with their optimised value) and a description of their purpose.

| Hyperparameter (=val) | Description |
| --- | --- |
| dl_nums_workers (=2) | To speed up the training process, we make use of the num_workers optional attribute of the DataLoader class.<br>The num_workers attribute tells the data loader instance how many sub-processes to use for data loading. By default, the num_workers value is set to zero, and a value of zero tells the loader to load the data inside the main process.<br>This means that the training process will work sequentially inside the main process. After a batch is used during the training process and another one is needed, we read the batch data from the disk.<br>Now, if we have a worker process, we can make use of the fact that our machine has multiple cores. This means that the next batch can already be loaded and ready to go by the time the main process is ready for another batch. This is where the speed up comes from. The batches are loaded using additional worker processes and are queued up in memory. |
| ims_per_batch (=2) | If we use 16 GPUs and IMS_PER_BATCH = 32, each GPU will see 2 images per batch. |
| gamma (=0.1) | The iteration number to decrease the learning rate by GAMMA. |
| backbone_freeze_at (=3) | Freeze the first several stages so they are not trained. There are 5 stages in ResNet. The first is a convolution, and the following stages are each group of residual blocks.<br>Freezing a layer prevents its weights from being modified. This technique is often used in transfer learning, where the base model (trained on some other dataset) is frozen. |
| warmup_iters (=120) | Warm-up steps are just a few updates with low learning rate before training. After this warm-up, the regular learning rate is used, (schedule) to train the model to convergence. The idea is that this helps the network to slowly adapt to the data intuitively.<br>If the data set is highly differentiated, it can suffer from a sort of "early over-fitting". If your shuffled data happens to include a cluster of related, strongly-featured observations, your model's initial training can skew badly toward those features - or worse, toward incidental features that are not truly related to the topic at all.<br>Warm-up is a way to reduce the primacy effect of the early training examples. Without it, one may need to run a few extra epochs to get the convergence desired, as the model un-trains those early superstitions.<br>Many models afford this as a command-line option. The learning rate is increased linearly over the warm-up period. If the target learning rate is $p$ and the warm-up period is $n$, then the first batch iteration uses $1p/n$ for its learning rate; the second uses $2p/n$, and so on: iteration $i$ uses $i * p/n$, until we hit the nominal rate at iteration $n$.<br>This means that the first iteration gets only $1/n$ of the primacy effect. This does a reasonable job of balancing that influence.<br>Note that the ramp-up is commonly on the order of one epoch – but is occasionally longer for particularly skewed data, or shorter for more homogeneous distributions. One may want to adjust, depending on how functionally extreme ones batches can become when the shuffling algorithm is applied to the training set. |
| momentum (=0.9) | Momentum in neural networks is a variant of the stochastic gradient descent. It replaces the gradient with a momentum which is an aggregate of gradients. Momentum can increase speed when the cost surface is highly non-spherical because it damps the size of the steps along with directions of high curvature thus yielding a larger effective learning rate along with the directions of low curvature. |
| batch_size_per_image (=1024) | The batch size defines the number of samples that will be propagated through the network. For instance, let's say you have 1050 training samples and you want to set up a batch_size equal to 100. The algorithm takes the first 100 samples (from 1st to 100th) from the training dataset and trains the network. Next, it takes the second 100 samples (from 101st to 200th) and trains the network again. We can keep doing this procedure until we have propagated all samples through of the network. Problem might happen with the last set of samples. In our example, we have used 1050 which is not divisible by 100 without remainder. The simplest solution is just to get the final 50 samples and train the network.<br>Advantages of using a batch size < number of all samples:<br>It requires less memory. Since you train the network using fewer samples, the overall training procedure requires less memory. That is especially important if you are not able to fit the whole dataset in your machine's memory. Typically networks train faster with mini-batches. That is because we update the weights after each propagation. In our example we have propagated 11 batches (10 of them had 100 samples and 1 had 50 samples) and after each of them we have updated our network's parameters. If we used all samples during propagation we would make only 1 update for the network's parameter.<br>Disadvantages of using a batch size < number of all samples:<br>The smaller the batch the less accurate the estimate of the gradient will be. |
| weight_decay (=0.001) | Weight decay is a regularization technique by adding a small penalty, usually the L2 norm of the weights (all the weights of the model), to the loss function. $\text{loss} = \text{loss} + \text{weight decay parameter} * \text{L2 norm of the weights}$.<br>Why do we use weight decay? To prevent overfitting. To keep the weights small and avoid exploding gradient. Because the L2 norm of the weights are added to the loss, each iteration of your network will try to optimize/minimize the model weights in addition to the loss. |
| base_lr (=0.0003389) | In machine learning and statistics, the learning rate is a tuning parameter in an optimization algorithm that determines the step size at each iteration while moving toward a minimum of a loss function. Since it influences to what extent newly acquired information overrides old information, it metaphorically represents the speed at which a machine learning model "learns". In the adaptive control literature, the learning rate is commonly referred to as gain.<br>In setting a learning rate, there is a trade-off between the rate of convergence and overshooting. While the descent direction is usually determined from the gradient of the loss function, the learning rate determines how big a step is taken in that direction. A too high learning rate will make the learning jump over minima but a too low learning rate will either take too long to converge or get stuck in an undesirable local minimum.<br>In order to achieve faster convergence, prevent oscillations and getting stuck in undesirable local minima the learning rate is often varied during training either in accordance with a learning rate schedule or by using an adaptive learning rate. |
| max_iter (=462) | An iteration describes the number of times a batch of data passed through the algorithm. In the case of neural networks, that means the forward pass and backward pass. So, every time you pass a batch of data through the neural network, you completed an iteration. When you increase the number of iterations, you get closer to the minima and the set of optimal parameters, and these are what improve performance. |

**Fig. S2.**
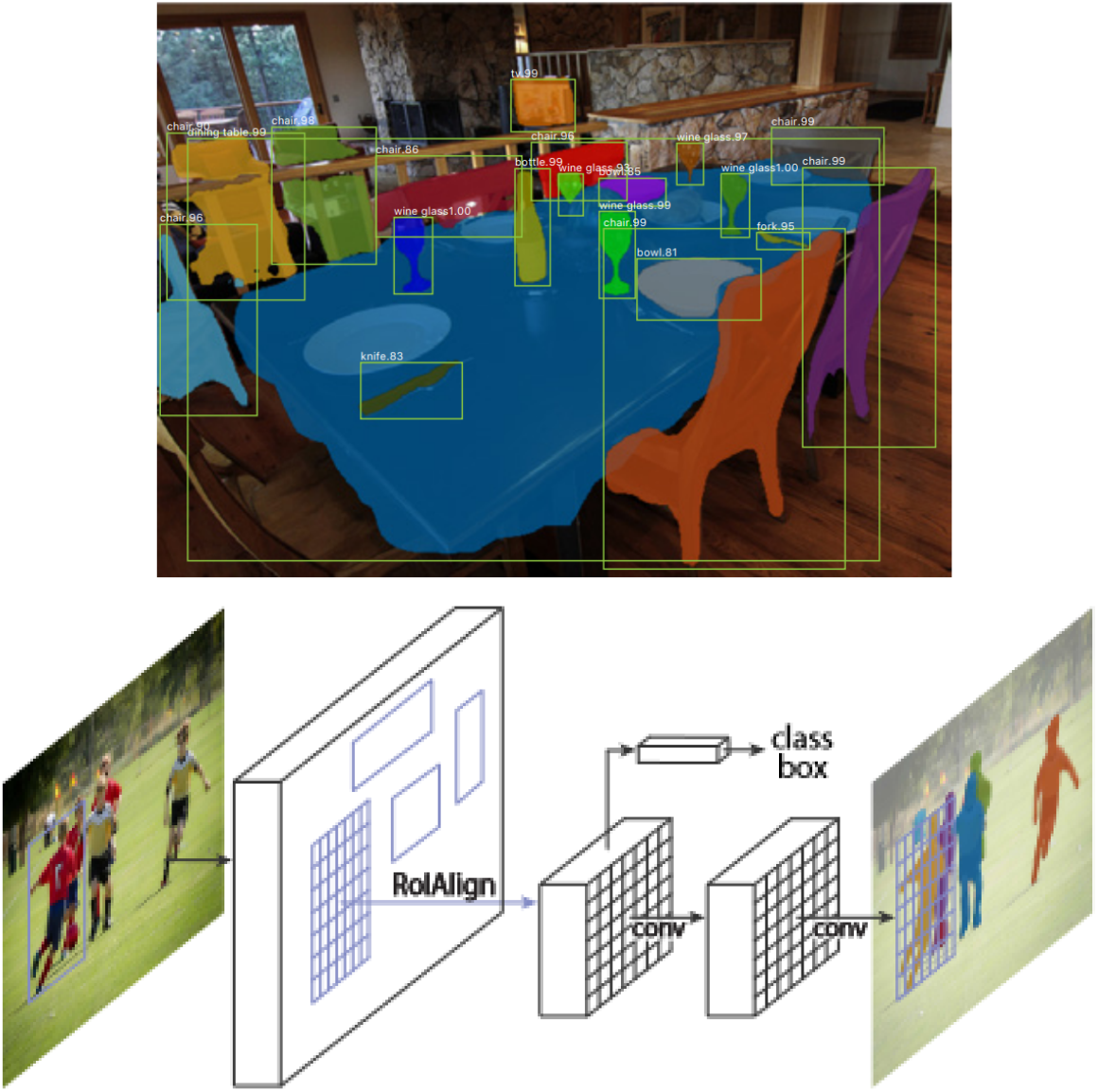
Illustration of the Mask R-CNN predictions and architecture from He et al. (40).

We found that while increasing the depth of the ResNet increased the training time of the algorithm, it improved accuracy scores. The batch size was limited by computing resources available. The number of iterations was the key metric to optimise, as we found that our model began to overfit the training data if trained for too many iterations. To determine the number of iterations, we plotted a graph (Fig. S6) of both the total training loss and the total validation loss to determine the minimum of the total validation loss, and hence the optimal number of iterations to train the model. As Mask R-CNN consists of three predictions, namely the class, bounding box and mask, the total loss is the sum of these three losses.

We see that after around 800 iterations, the model started to overfit the training data, at the expense of performance of the validation dataset. As such we determined to stop the training of Mask R-CNN at 800 iterations, and then used the saved weights after 800 iterations to predict tree crowns in Sepilok.

## Supplementary Note 5: Evaluation metrics

Two key metrics are used in this work to evaluate the performance of the model: the AP50 score, and the F_1_ Score. AP50 relates to the area under the precision-recall curve (AUC-PR) evaluated at a particular threshold for the Intersection over Union, in our case 0.5. A high value for AP50 means that we are seeing both high precision and recall of the model at our particular IoU cutoff. The IoU cutoff was selected as 0.5, on the basis of previous studies evaluating the accuracy of individual tree delineation methods. In rare cases where more than one automatic delineation had an *IoU* of greater than 0.5 with a manual crown, then the automatic delineation with the greatest *IoU* was considered a true positive and the others were classed as false positives. Visual examples of different *IoU* values are given in Fig. 1.

**Fig. S3.**
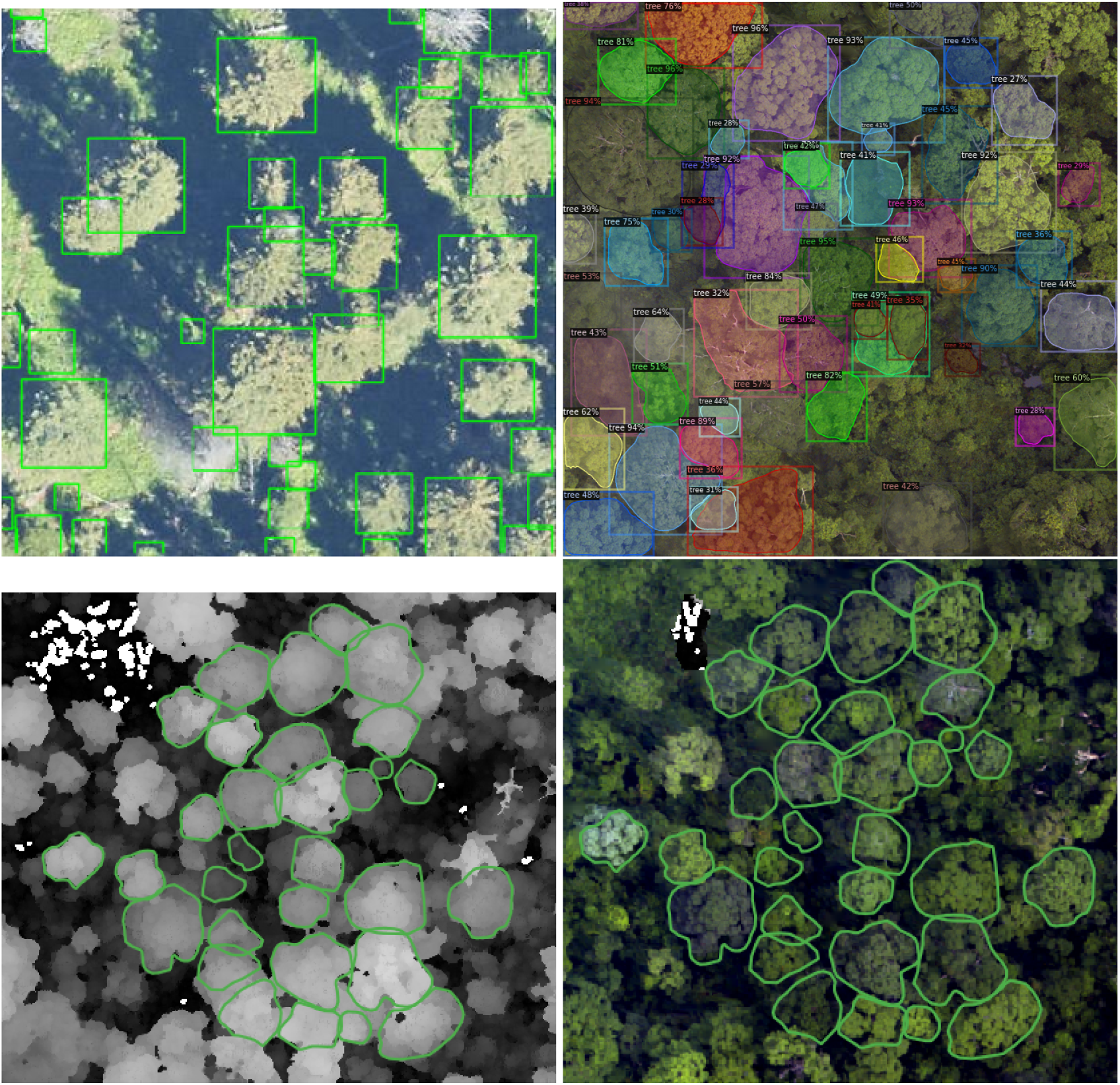
(A) Crown bounding boxes predicted by DeepForest (Weinstein et al., 2019), and (B) crowns predicted by Detectree2. The coIoUrs in plot B merely distinguish predicted trees. A comparison of manually delineated crowns, overlaid on lidar (C) and RGB (D).

**Fig. S4.**
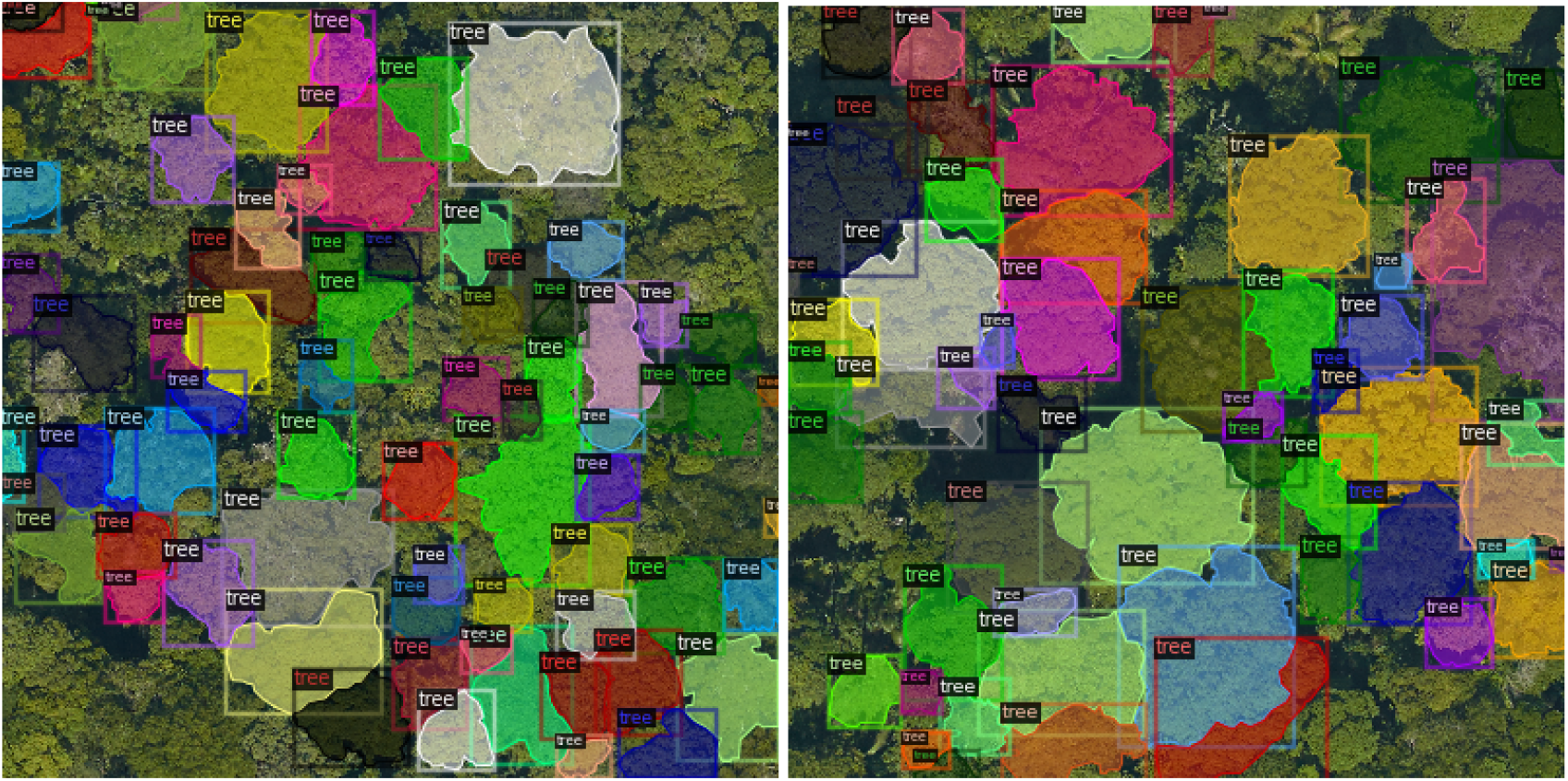
Examples of training data provided to Mask R-CNN. The different coIoUrs help to distinguish between trees.

**Fig. S5.**
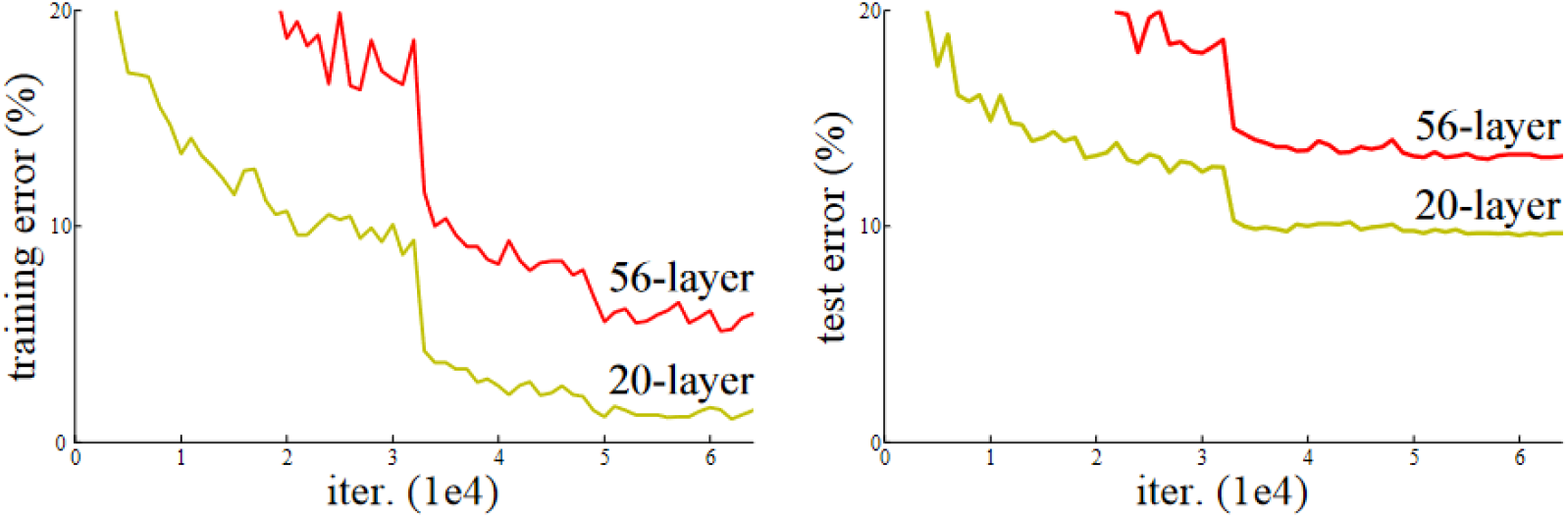
These plots are taken from He et al., (2016) and they illustrate that deeper neural networks do not necessarily learn as well as shallower neural networks.

**Fig. S6.**
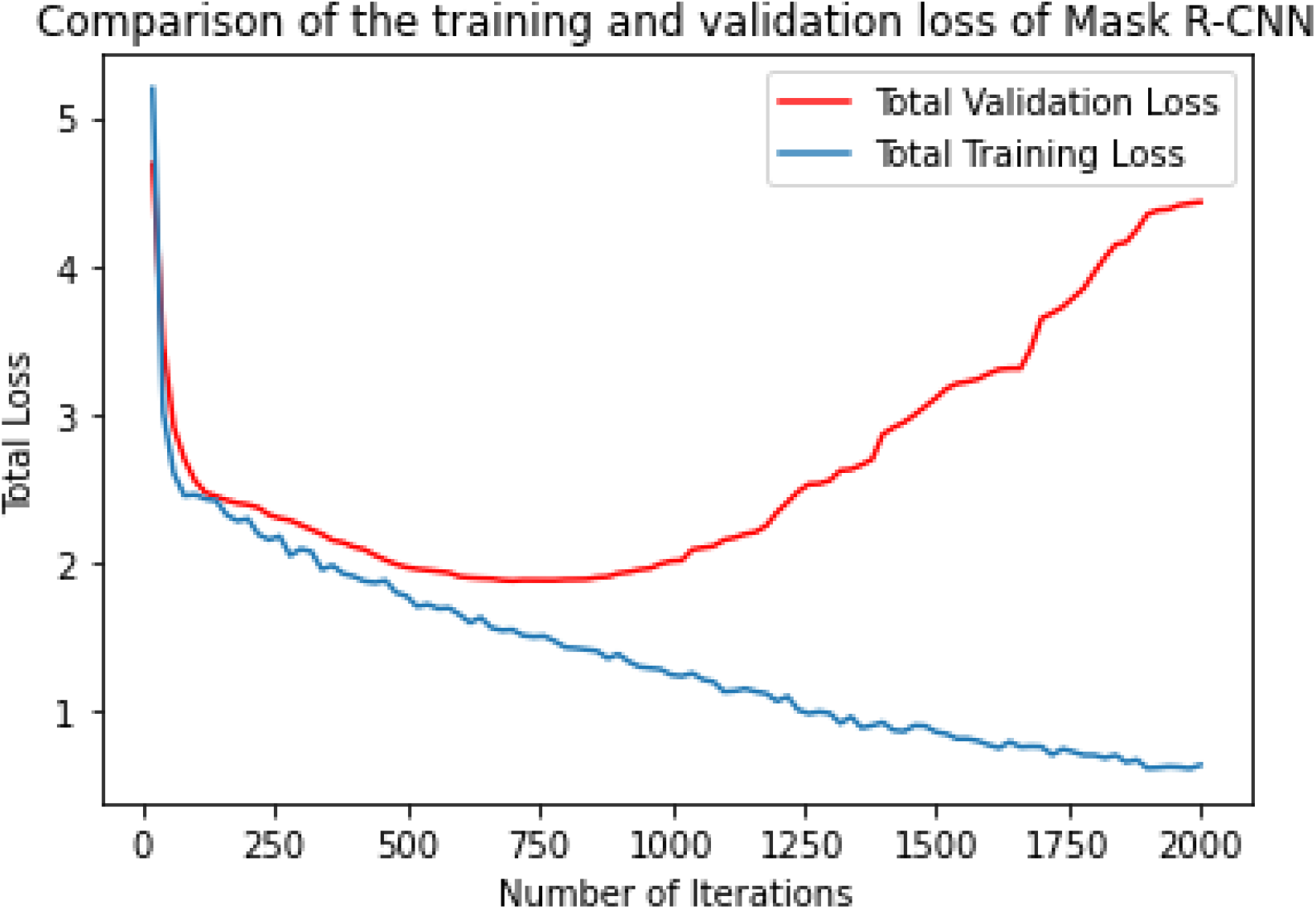
The total training and validation loss of Mask R-CNN as the model trained. Both total training and validation loss were calculated every 20 iterations.

## Supplementary Note 6: Maps of predictions

**Fig. S7.**
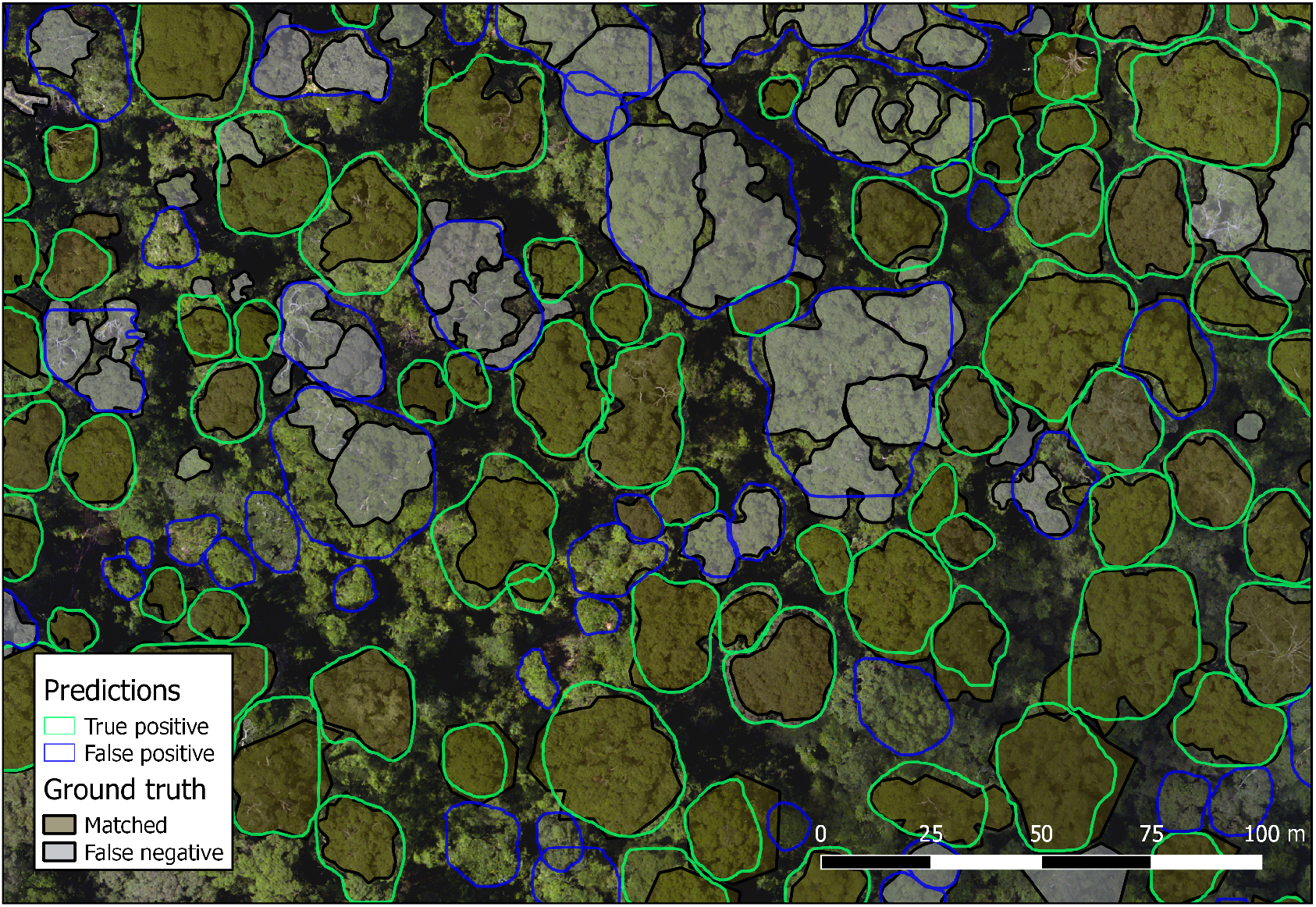
Example delineation results at Danum

**Fig. S8.**
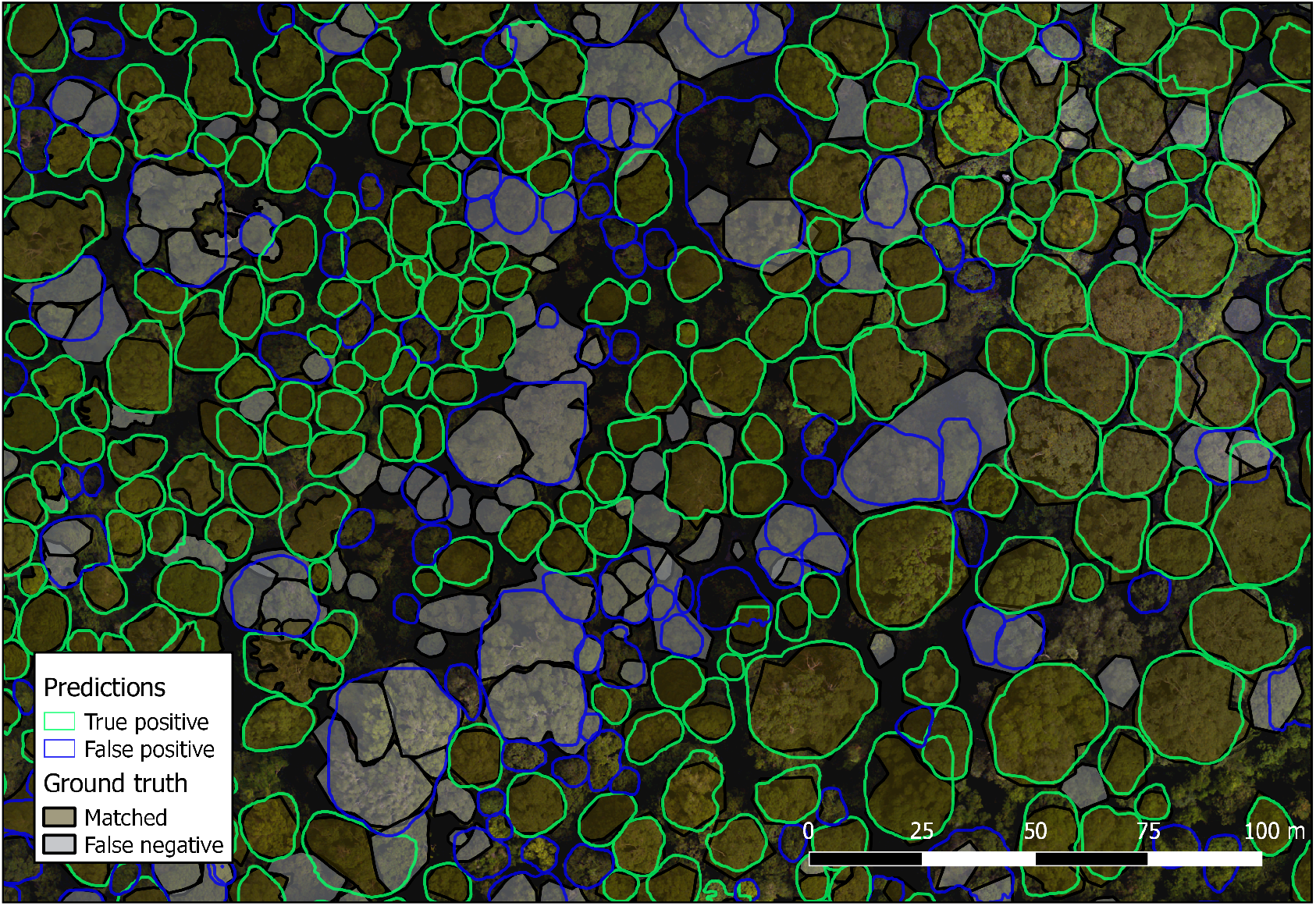
Example delineation results at Sepilok West

**Fig. S9.**
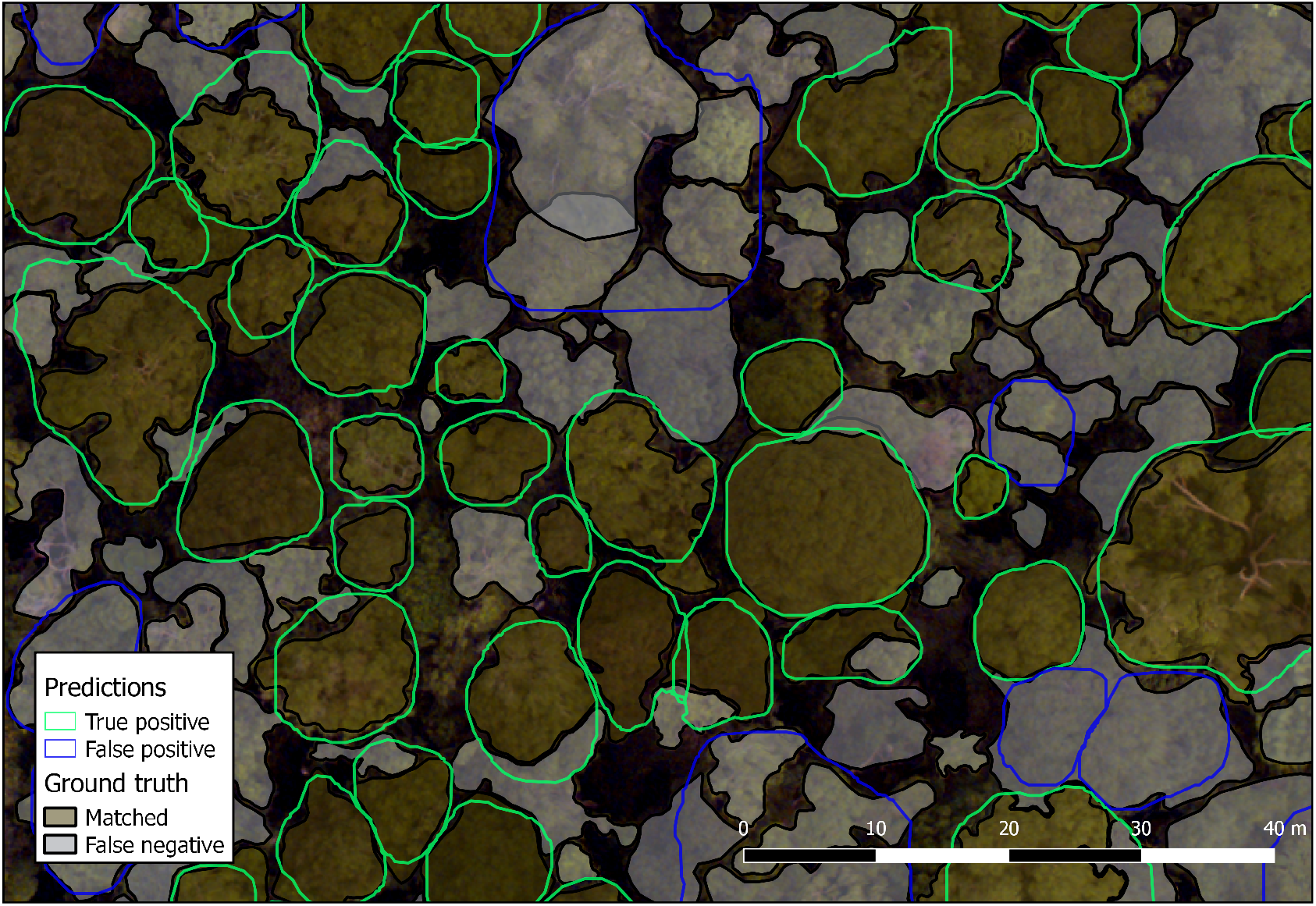
Example delineation results at Sepilok West

**Fig. S10.**
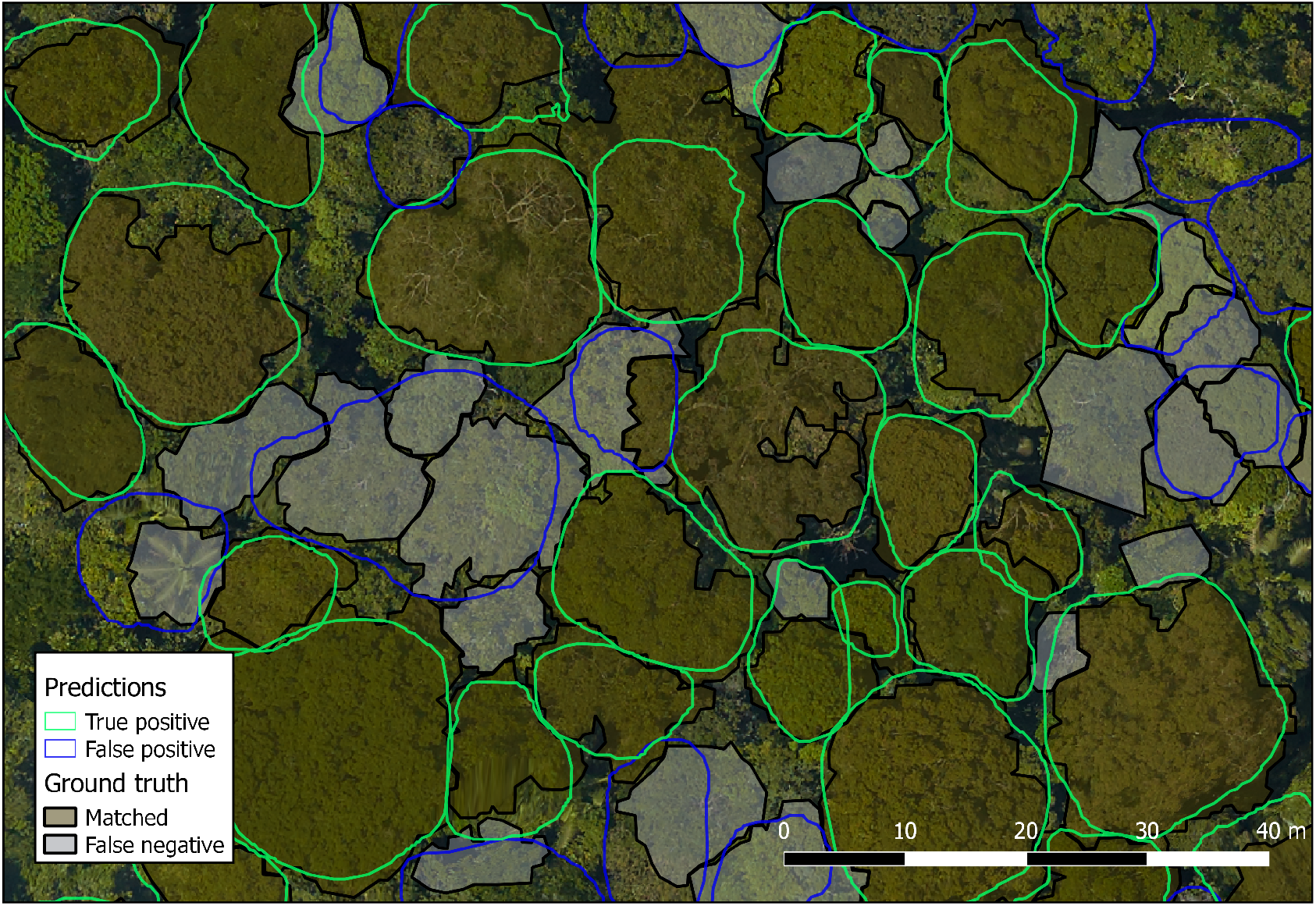
Example delineation results at Paracou

## Supplementary Note 7: Sensitivity to image resolution

We performed a preliminary analysis on how sensitive the delineation accuracy is to changes in the image resolution. We then carried out another test, where we trained our model on the highest resolutions available to us (8 and 10 cm), before testing it on lower resolutions to determine the model’s sensitivity to resolution. The lower resolutions were chosen as they are typical resolutions of modern, high-resolution satellite imagery. The final test was to train and test the model on low resolutions to determine if this workflow would increase the skill of the model on lower resolutions. As seen in Fig. S11, illustrate that our model trained on high resolution (0.1m) imagery is most successful when predicting on similar resolutions, and its performance degrades for resolutions an order of magnitude greater (1 m and 2 m). However, we illustrate that when the model is trained on similar resolutions to those it is tested on, performance is largely maintained. It is only on resolutions greater than 1 m that the performance degrades significantly.

**Fig. S11.**
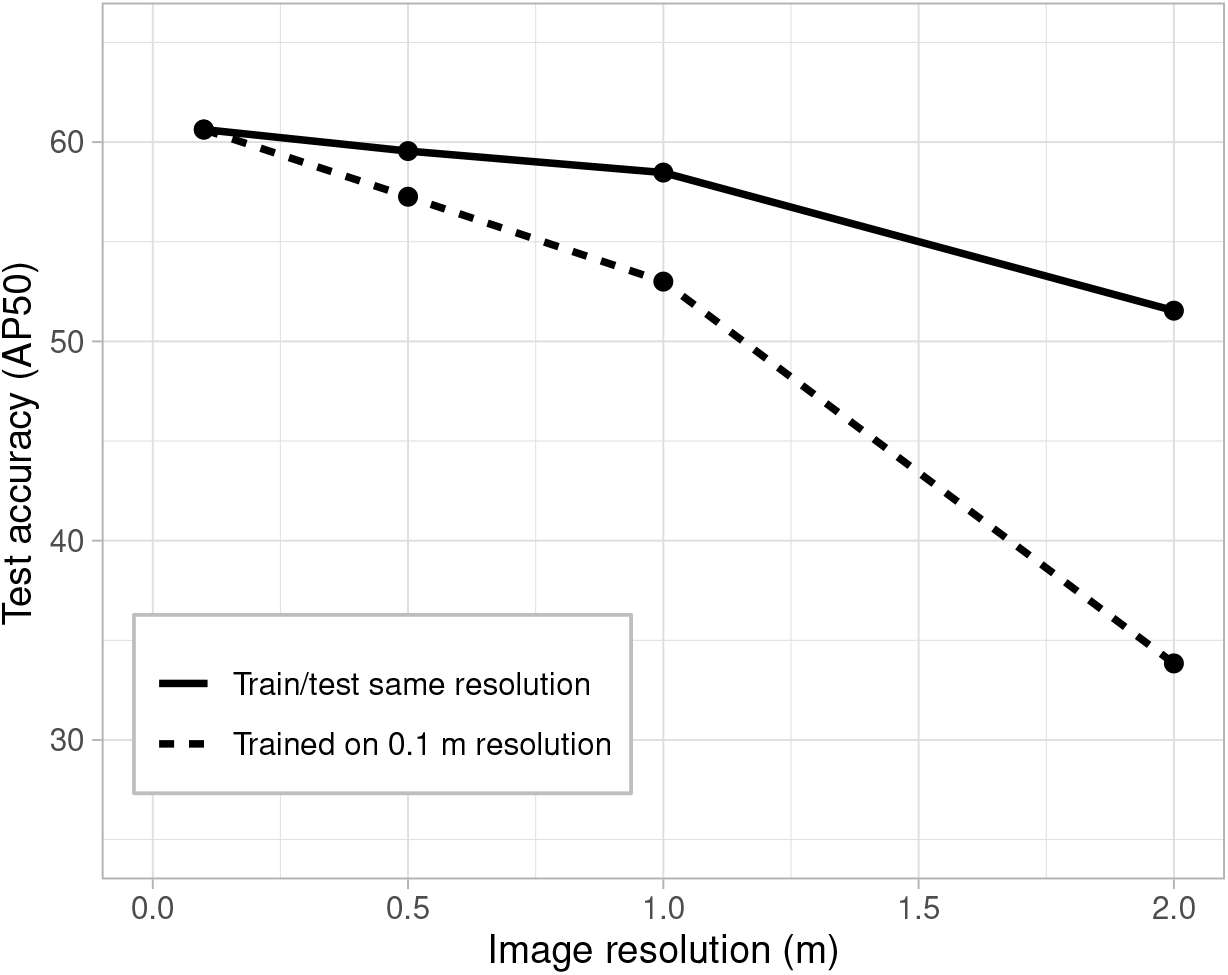
The sensitivity of the accuracy of the segmentations to the resolution of images used in training and testing.

## Supplementary Note 8: Growth and mortality details

Growth and mortality rates were estimated from the CHMs of the repeat lidar data (taken at all sites as detailed in Table 1). Tree height was defined by the median value of the CHM within the overlayed crown polygon. To calculate growth rate (in m / yr), we evaluated the difference in height of individual trees in the two years of measurements, and divided by the time between the measurements (assuming an approximately constant growth rate).

Mortality estimates were derived by fitting a robust least squares regression to the change in tree height against the original tree height. Mortality events were defined as a drop in height of 3 standard deviations or more from this fit. The rate of mortality (%/yr) was calculated as the percentage of detected trees that died divided by the time between scans. The tree mortality rates were grouped into height bins (depending on the original height of each tree). A drop of three standard deviations is strict but it is possible that drops could result from non-mortality events such as snapping or other wind damage. Uncertainty estimates were determined by bootstrapping (repeatedly sampling from the complete set of predicted crowns).

**Table S2.** The coefficients and intercepts for the robust least squares fit between original tree height and the change in tree height.

| Site | Slope | Intercept |
| --- | --- | --- |
| Danum | $-0.110 \pm 0.006$ | $1.86 \pm 0.26$ |
| Paracou | $-0.028 \pm 0.0003$ | $1.06 \pm 0.01$ |
| Sepilok East | $-0.060 \pm 0.004$ | $1.67 \pm 0.10$ |
| Sepilok West | $-0.072 \pm 0.003$ | $2.72 \pm 0.12$ |

Plots analogous to Fig. 5(A) for the other sites are given in Fig. S12. These plots illustrate the different characteristics of each forest site.

**Fig. S12.**
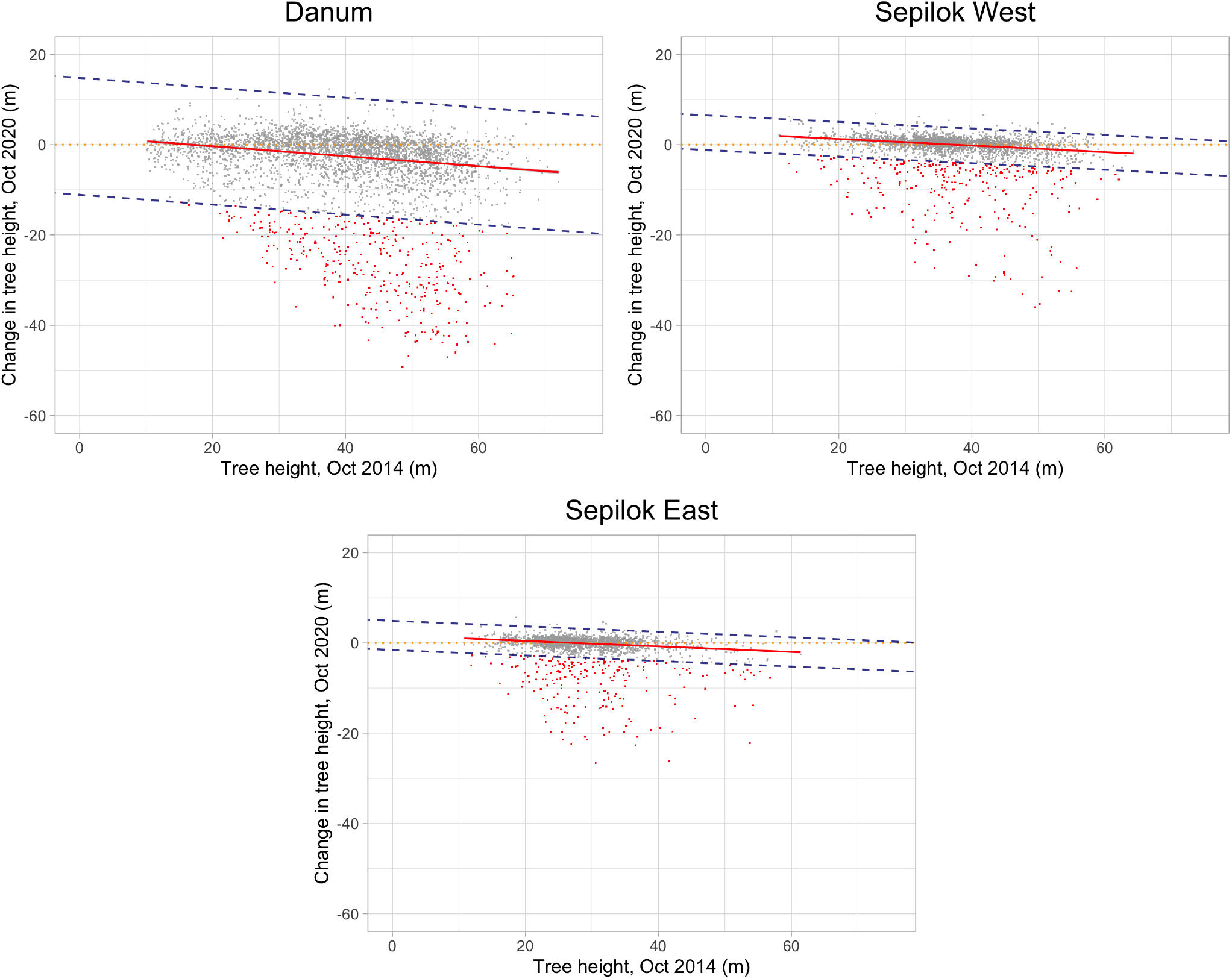
shows the robust least squares fit for change in height and tree height for Sabah (Danum, Sepilok West and Sepilok East). The dashed lines indicate three standard deviations either side of the best fit and red points below the lower bound indicate likely mortality events.

## Footnotes

1 https://github.com/PatBall1/Detectree2

2 https://github.com/facebookresearch/detectron2/blob/main/MODEL_ZOO.md

3 https://github.com/facebookresearch/detectron2/blob/main/configs/COCO-Detection/faster_rcnn_R_101_FPN_3x.yaml

4 https://wandb.ai/detectree/tune/sweeps/

5 https://github.com/PatBall1/Detectree2

6 See https://github.com/PatBall1/Detectree2

7 https://www.image-net.org/

